# Neural mechanisms of credit assignment for delayed outcomes during contingent learning

**DOI:** 10.1101/2024.08.06.606895

**Authors:** Phillip P. Witkowski, Lindsay Rondot, Zeb Kurth-Nelson, Mona M. Garvert, Raymond J Dolan, Timothy E. J. Behrens, Erie D. Boorman

## Abstract

Adaptive behavior in complex environments critically relies on the ability to appropriately link specific choices or actions to their outcomes. However, the neural mechanisms that support the ability to credit only those past choices believed to have caused the observed outcomes remain unclear. Here, we leverage multivariate pattern analyses of functional magnetic resonance imaging (fMRI) data and an adaptive learning task to shed light on the underlying neural mechanisms of such specific credit assignment. We find that the lateral orbitofrontal cortex (lOFC) and hippocampus (HC) code for the causal choice identity when credit needs to be assigned for choices that are separated from outcomes by a long delay, even when this delayed transition is punctuated by interim decisions. Further, we show when interim decisions must be made, learning is additionally supported by lateral frontopolar cortex (lFPC). Our results indicate that lFPC holds previous causal choices in a “pending” state until a relevant outcome is observed, and the fidelity of these representations predicts the fidelity of subsequent causal choice representations in lOFC and HC during credit assignment. Together, these results highlight the importance of the timely reinstatement of specific causes in lOFC and HC in learning choice-outcome relationships when delays and choices intervene, a critical component of real-world learning and decision making.

## Introduction

Humans and animals have a remarkable ability to navigate complex environments and infer the likely state of the world from observed phenomena. Such adaptive behavior requires the ability to learn about causal relationships between one’s choices and subsequent outcomes. A key challenge for learning systems in the brain arises when a task involves temporal delays between choices and their outcomes. Cooking is one such task in which many decisions may be made about how to adjust the flavor profile of a dish, but the resultant outcomes of these choices typically will not be evaluated until sitting down to eat. Moreover, cooking often requires juggling multiple sub- tasks simultaneously, meaning that interim decisions need to be performed in between adding an ingredient and observing its effect on the dish’s flavor. In such cases, discerning the causal relationship between a particular choice and possible outcomes is nontrivial. While this ability to link choices and outcomes is critical to success in real-world tasks, little is known about *how* these links are forged at the neural level.

A large body of pioneering work focusing on the role of the lateral orbitofrontal cortex (lOFC) has highlighted the importance of this region in contingent learning (Gardner & Schoenbaum, 2021; Murray & Rudebeck, 2018; Rushworth et al., 2011). Recent studies in multiple species have emphasized a special role for lOFC in leveraging task knowledge for credit assignment, linking specific reinforcement outcomes to specific past choices (Boorman et al., 2013; Jocham et al., 2016; Lamba et al., 2023; Stalnaker et al., 2015; Sutton & Barto, 2014; Walton et al., 2010). In one key study, lesions to the macaque lOFC, impaired the ability of animals to use a model of the task structure in order to track the contingency between specific choices and outcomes they caused, with credit erroneously spreading to non-causal choices (Walton et al., 2010). These results suggest that lOFC is required for using a model of the task structure to form, or update, an association between specific choices and outcomes. Such findings were subsequently replicated and extended in both rats and humans (Costa et al., 2023; Noonan et al., 2017). Other studies in humans have shown that outcome-related blood oxygen-level-dependent (BOLD) activity in lOFC is specific to contingent, but not non-contingent, reward observations (Jocham et al., 2016), and the magnitude of activity reflects the degree to which credit for an outcome is assigned (Boorman et al., 2013, 2016). Collectively, these findings suggest that computations within the lOFC are critical to credit assignment; however, little is known about the mechanisms by which the lOFC supports assigning credit for outcomes to specific causes.

One possible mechanism by which the brain assigns credit when reinforcement is delayed is by reinstating a representation of the causal choice at the time of feedback. In principle, this could enable the choice representation to be associated with the online encoding of the outcome, potentially via changes in synaptic plasticity between co-active neuronal ensembles. Such coding of past choices specifically at the time of feedback has been identified in macaque lOFC neuronal ensembles, albeit in the absence of any task requirement for contingent learning (Tsujimoto et al., 2009). Likewise, altered dopaminergic prediction error responses in lOFC-lesioned rats were elegantly accounted for by a computational model that incorporates a loss of internal representations of an outcome-linked choice, leading to misattributing value across states (Takahashi et al., 2011). Information about previous choices is also found in regions to which the lOFC shares reciprocal connectivity, particularly the hippocampus (HC) (Barbas & Blatt, 1995; Wikenheiser & Schoenbaum, 2016). A largely separate literature focusing on HC has shown reinstatement of neural activity patterns previously elicited by a stimulus both at the time of choice and reward in sensory pre-conditioning paradigms (Barron et al., 2020; Kurth-Nelson et al., 2015; Wimmer & Shohamy, 2012), and likewise during associative inference and integration (Koster et al., 2018; Park et al., 2020; Zeithamova et al., 2012). Such hippocampal reinstatement of stimulus identity representations might be expected to support lOFC coding of relevant past choices for credit assignment, particularly following lengthier delays (Foerde & Shohamy, 2011; Shohamy et al., 2009; Wang et al., 2020).

In complex tasks where subsequent decisions intervene on the transitions between choices and resultant outcomes, the neural regions supporting credit assignment may extend to encompass regions that also support maintaining information about causal choices pending their resultant outcome. This would allow learning systems to precisely assign credit to causal choices by bridging over interim decisions that may otherwise be inappropriately linked to the observed outcome. A key region for maintaining such “pending” information is the lateral frontal pole (lFPC), which has been implicated in maintaining information about prospective actions or cognitive processes that must be delayed and performed in the future (Burgess et al., 2007, 2011, 2022). Other research has shown that lFPC activity reflects the reliability of pending alternative task sets (Donoso et al., 2014; Koechlin et al., 2003; Koechlin & Hyafil, 2007), and that it tracks evidence favoring adapting behavior to specific counterfactual alternatives, and directed exploratory choices, in the future (Badre et al., 2012; Boorman et al., 2009, 2011; Zajkowski et al., 2017). On this basis, we hypothesized that the lFPC would play a critical role in maintaining information about previous choices that will be needed for future credit assignment during interim decisions.

In the current study, we test these hypotheses using a learning task in which participants must track contingencies between specific choices and outcomes under conditions where choice-outcome transitions are direct following a delay, or indirect and involve an intervening decision. We show that in both conditions, the lOFC and HC reinstate representations of causal choices at the time of feedback. In the indirect condition, this information is critically dependent on representations of the causal choice maintained in a “pending state” in lFPC, which predict subsequent reinstatement in lOFC and HC. Finally, we show that lOFC and HC code task-independent stimulus identity representations during feedback, suggesting a link between coding of a state’s identity and precise credit assignment.

## Results

### Learning task with direct and indirect choice-outcome transitions

Participants completed a learning task in which they chose between two abstract shapes to obtain one of two distinct outcomes (gift cards to locally available stores rated to be approximately equally desirable). Each shape had a certain probability of leading to one gift card and the inverse probability of leading to the other. These probabilities drifted over time but could be tracked based on the recent choice-outcome observations made in each trial (see Fig. S1 for probability trajectories and Bayesian model fitting). Participants were informed of how many points each gift card would yield on each trial by colored numbers on the top of the screen, and that these points changed randomly from one trial to the next (Fig. 1A). They were further told that at the end of the experiment one trial would be selected at random to count “for real”. That is, they would receive the gift card obtained on that trial with a value proportional to the number of points won. Thus, participants were incentivized to maximize their potential winnings on every trial by accurately tracking the probability that each shape would lead to each outcome, but not the history of reward amounts.

**Figure 1.**
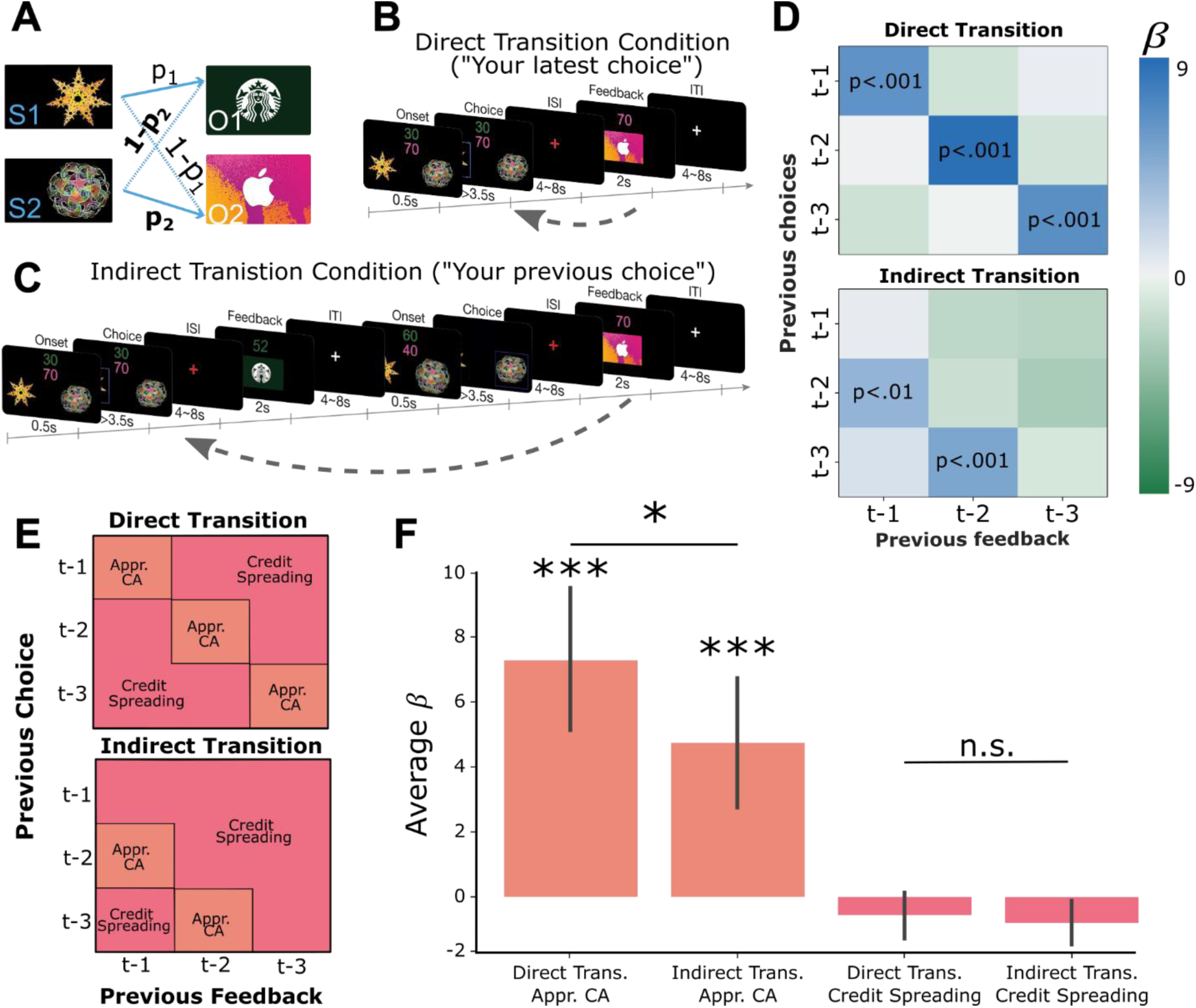
A) Two abstract shapes were probabilistically related to each of two outcome identities by independent transition probabilities p_1_ and p_2_. B) Schematic of the direct transition condition. Participants chose one of the two shapes on each trial based on two pieces of information: their estimates of the probability that each would lead to either outcome identity (gift cards) and the randomly generated number of points they could potentially win if that outcome was obtained. The color of each number indicated the identity of the outcome on which that number of points could be won. In the example, green indicates the number of points for the Starbucks gift card, while pink indicates the number of points for iTunes. Next, participants observed the outcome of their choice (the gift card and amount) after a delay. C) Schematic of the indirect transition condition. Same as (B) except that after participants made their choice they transitioned into another independent decision. After this second decision was made, participants observed the outcome of their first decision. D) Results of logistic regression analysis predicting the current choice based on previously observed choice-outcome relationships. Each cell represents the combination of a previously observed choice with an observed outcome. The color of each cell shows the value of beta estimates for each combination of previous choice and observed outcome, averaged across participants. Positive values indicate that the choice-outcome pair predicted choosing the same shape again when that shape previously led to the currently desired outcome. E) Theoretical decomposition of the matrix in (D) into groups of cells which reflect “appropriate credit assignment” given the task structure (orange) and “credit spreading” (pink). F) Mean (±SEM) of beta coefficients for specific choice-outcome combinations averaged across the groupings of cells shown in E for each condition.

The task had two conditions which proceeded in a blocked fashion. In the “direct transition” condition, participants saw the outcome of a choice after a delay period (Fig. 1B). In the “indirect transition” condition, participants did not see the outcome of their choice until *after* another choice had been made, requiring them to delay assigning credit to the initial choice until the appropriate outcome was observed (Fig. 1C). Participants were instructed about which condition they were in with a screen displaying “Your latest choice” in the direct transition condition, and “Your previous choice” in the indirect condition. Finally, at the beginning of each block participants viewed each of the two abstract shapes and two outcome stimuli in a random order, without making decisions or observing outcomes. This “template” block allowed us to measure neural responses to stimuli independently of the learning task.

#### Predicting current choice based on previous choice-outcome relationships

To test whether participants were using the structure of each condition to appropriately assign credit to causal choices, we performed a multiple logistic regression analysis testing the influence of previous choice-outcome combinations on the current choice. For each participant, independently in each condition, we constructed a GLM that predicted the current choice as a function of nine different combinations of previous choices and outcomes (Eq.1). For example, the first regressor predicted the current choice based on the previous choice and the previous outcome (trial *t-1*). These values were coded as 1 if the past choice led to the currently desired outcome, assumed to be the outcome with the largest monetary point value on the current trial, and -1 if it did not (results were virtually identical if we used the participant-specific indifference point (α) to define the desired outcome instead (see Eq. 9)). The second regressor predicted the current choice based on the previous choice (*t-1*) and the outcome received two trials in the past (*t-2*), and so on for all nine combinations of previous choices and outcomes covering the previous three trials.

In the direct transition condition, we observed significant positive effects along the diagonal of the matrix (*choice*_*t*−1_ ∗ *outcome*_*t*−1_ : β = 6.09, *t*(19) = 4.81, *p <* 0.001; *choice*_*t*−2_ ∗ *outcome*_*t*−2_: β = 8.78, *t*(19) = 5.41, *p <* 0.001; *choice*_*t*−3_ ∗ *outcome*_*t*−3_, β = 6.76, *t*(19) = 4.16, *p <*0.001; Fig. 1D), indicating that participants assigned credit for each outcome to the choice made in same trial. In the indirect transition condition, current choices were significantly predicted by the most recently observed outcomes combined with choices made in the trial previous to those outcomes (*choice*_*t*−2_ ∗ *outcome*_*t*−1_: β = 4.20, *t*(19) = 2.92, *p <*0.01; *choice*_*t*−3_ ∗ *outcome*_*t*−2_: β = 5.07, *t*(19) = 4.75, *p <*0.001). Furthermore, the mean of the β-values which reflect appropriate credit assignment in each condition were significantly higher than the mean β-values which represented credit spreading (direct transition condition: *t*(19) = 5.39, *p* < 0.001, indirect transition condition: *t*(19) = 4.34, *p*<0.001; Fig. 1E and F). Follow-up analysis showed that participants’ choices in each trial integrated expectations about the probability of receiving a particular outcome and its magnitude and did not rely on estimates of a cached option value (Fig. S1). These results show that participants used the appropriate task-structure when assigning credit for observed outcomes in each condition.

Next, we compared the relative precision of credit assignment between our behavioral conditions, where we predicted credit assignment would be less precise in the indirect transition condition compared to direct transition condition, owing to additional task complexity. We found that β-values representing appropriate credit assignment in the direct transition condition were higher than those in the indirect transition condition (*t*(19) = 1.81, *p* <0.05). However, β-values in cells that represent credit spreading in the direct transition condition were not significantly lower than those in the indirect transition condition (*t*(19) = 1.11, *p*=.14). These results indicate that credit assignment was less precise in the indirect transition condition compared to the direct transition condition, despite each being appropriate for the respective task structure overall.

#### Causal choice codes are reinstated in lOFC and HC when viewing the outcome of choices

For the direct feedback condition, our main hypothesis was that lOFC codes for the specific causal choice when participants view the outcome of their choice. We also reasoned that, due to the delay between choice and feedback, this lOFC choice code would be supported by choice reinstatement in the interconnected HC (Barbas & Blatt, 1995; Wimmer & Shohamy, 2012). We tested this hypothesis by training a linear support vector machine (SVM) to distinguish BOLD activity patterns at the time of feedback based on the previously chosen shape, cross-validated across scanning runs (see Methods for details on decoding procedure). We used a searchlight analysis within *a priori* defined ROIs for lOFC and HC to estimate decoding accuracy for each voxel within the ROI (Kriegeskorte et al, 2008).

We found evidence for choice decoding in the predicted network of regions. Specifically, we found significant and marginally significant decoding of the causal choice in left ([x,y,z] = [- 26, 42, -8], *t*(19) = 4.22, *pTFCE* <0.05 ROI-corrected using threshold-free cluster enhancement (TFCE) correction (Smith & Nichols, 2009)) and right ([x,y,z] = [24, 46, -8], *t*(19) = 3.45, *pTFCE* = 0.081 ROI-corrected]) lOFC, respectively (Fig. 2A). A similar pattern was also apparent in the HC, where right HC showed significant decoding ([x,y,z] = [36, -20, -16], *t*(19) = 4.02, *pTFCE* <0.05 ROI-corrected]), while left HC showed a marginal effect ([x,y,z] = [-22, -10, -24], *t*(19) = 2.86, *pTFCE* = 0.080 ROI-corrected]). Together, these results show that the lOFC and HC represent the causal choice at the time when credit is assigned in the direct condition of our task.

**Figure 2.**
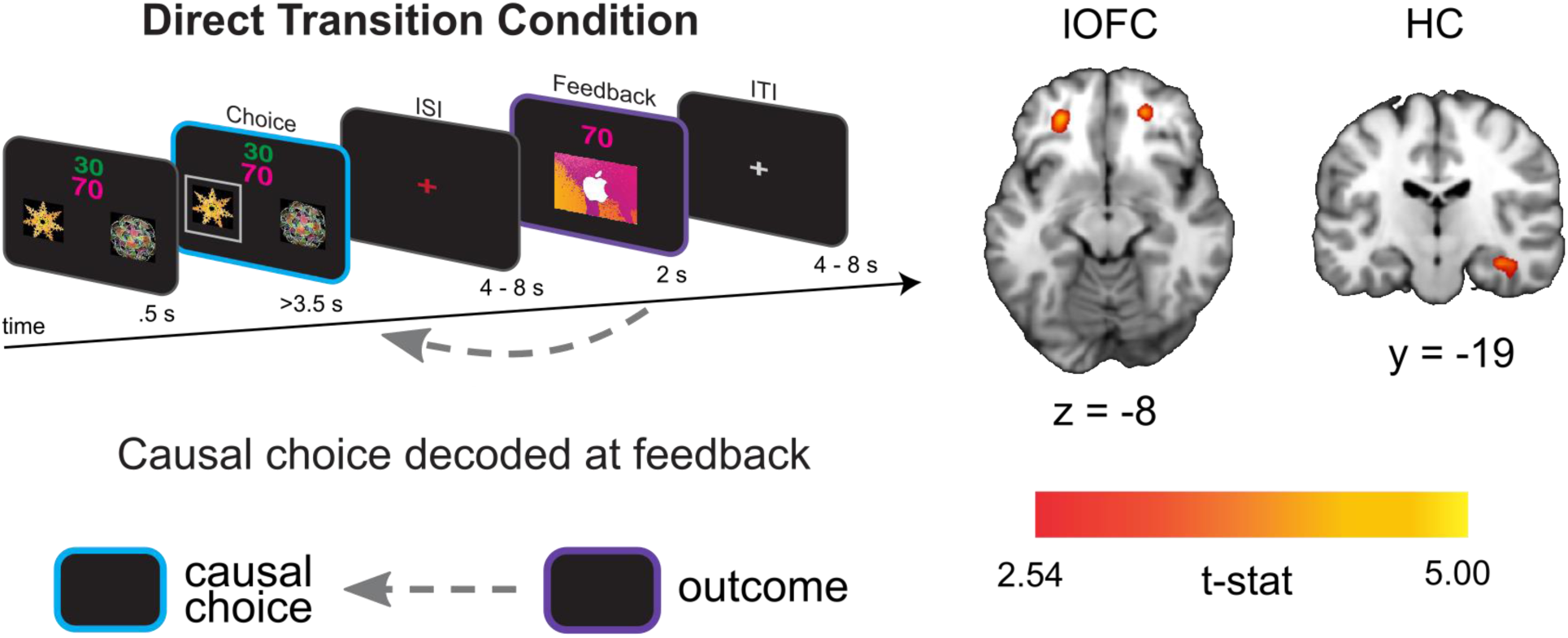
Left side shows the analysis scheme for decoding representations of the causal choice at feedback in the direct transition condition. An SVM decoder was used to differentiate trials at the time of the outcome (purple) based on the causal choice selected during the “choice period” (cyan). The right side shows axial and coronal slices through a t-statistic map showing significant decoding in OFC and HC during feedback. For illustration, all maps are displayed at threshold of t(19) = 2.54, p<0.01 uncorrected. All effects survive small volume correction in a priori defined anatomical ROIs.

#### Pending item representations in lFPC during indirect transitions predict credit assignment in lOFC

The indirect transition condition allowed us to test whether similar reinstatement mechanisms, as described above, support credit assignment when choice-outcome transitions are punctuated by interim decisions. We anticipated that the structure of the indirect transition condition would render credit assignment more difficult compared to the direct transition condition; a prediction borne out by our behavioral analysis of learning (Fig. 1F). Repeating the causal choice decoding analysis on this condition did not reveal a significant effect in any *a priori* defined ROI (all *pTFCE* >0.05 ROI corrected), nor did we find significant decoding elsewhere in the brain (all *pTFCE* >0.05 whole brain corrected). However, a key attribute of this condition is that causal choices must be held in a pending state during interim choices until a prospective outcome is observed. Thus, we reasoned that the fidelity of credit assignment at the time of feedback would be intimately related to the fidelity with which representations were maintained during the interim decision.

Following previous work suggesting that prospective representations of to-be-completed tasks are supported by lFPC (Burgess et al., 2011; Koechlin & Hyafil, 2007), we predicted that lFPC would hold causal choices in a “pending state” when credit assignment needs to be deferred until the resulting outcome is observed. To test this hypothesis, we used a linear SVM to classify neural activity at the time of feedback based on the immediately preceding choice. Note that in this condition the immediately preceding choice is *not* the cause of the currently observed outcome, but is the cause of the outcome for which credit will be assigned in the next trial. We call this the “pending causal choice”. Our analysis revealed a cluster of voxels specifically within the right lFPC ([x,y,z] = [28, 54, 8], *t*(19) = 3.74, *pTFCE* <0.05 ROI-corrected; left hemisphere all *pTFCE* > 0.1, Fig. 3A), consistent with right lFPC coding for the pending causal choice at feedback time, precisely when the outcome of the prior choice causal choice needed to be evaluated.

**Figure 3.**
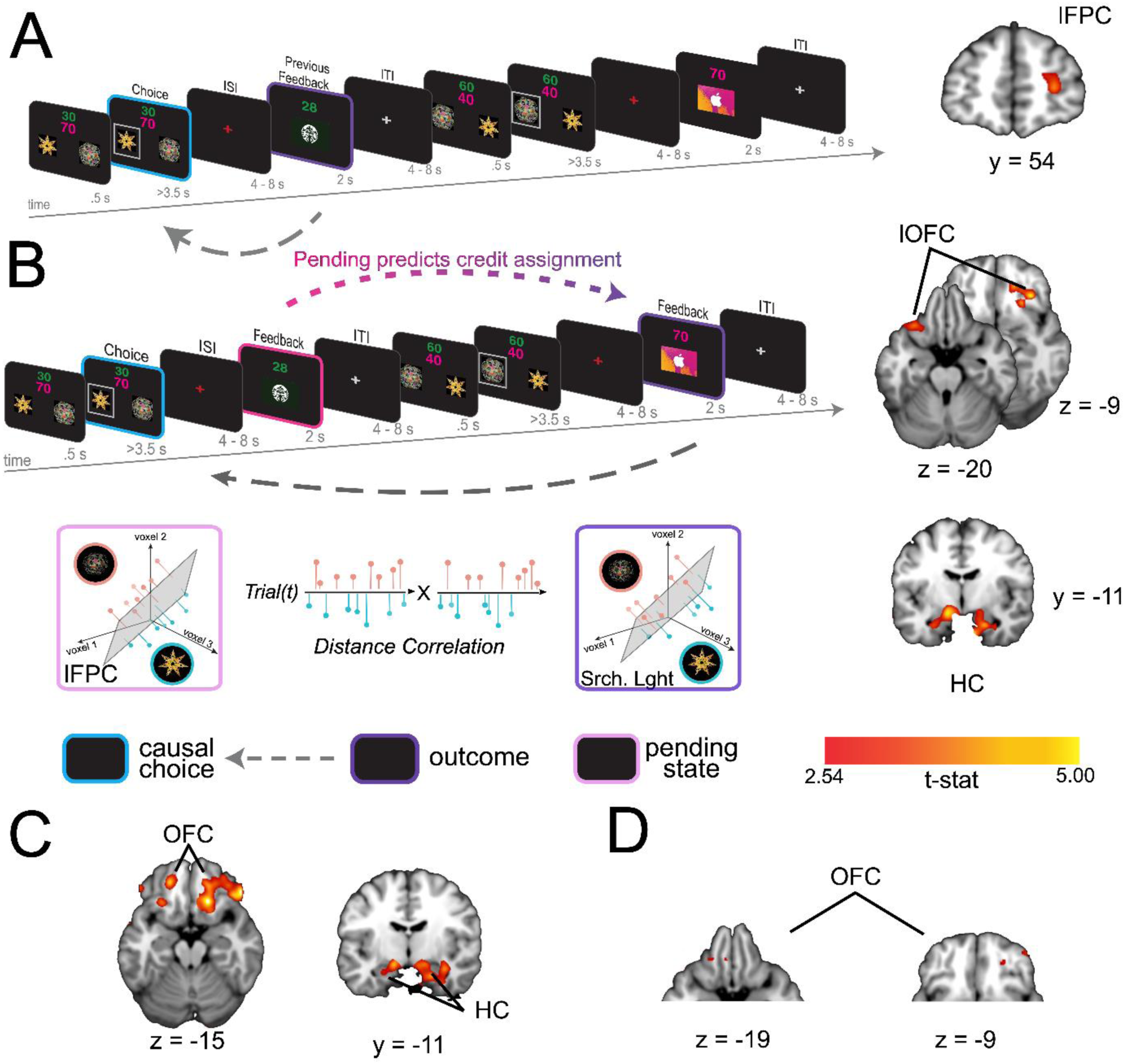
A) Left side shows the analysis scheme for decoding information about the causal choice in “pending state” (pink) in the indirect transition condition. We decoded information about the previous choice during the feedback period, during which the causal choice should be “pending” credit assignment in the next trial. The image on the right shows a coronal slice through a t- statistic map, showing significant decoding in lFPC. B) The analysis scheme for the information connectivity analysis which uses the trial-by-trial fidelity of causal choice representations in the “pending state” (pink) to predict the fidelity of these same choices when the outcome is observed (purple). The right side shows axial and coronal slices of a t-statistic map showing effects in lOFC and HC. All maps are displayed using the same conventions as Fig. 2 and all effects survive small volume correction in a priori defined anatomical ROIs. C) Axial (left) and coronal (right) slices through a t-statistic map showing the results of a control analysis in which we test the proportion of correct classifications of causal choice information in OFC and HPC at the time of the outcome for trials in which the lFPC showed correct classification for the causal choice during pending trials. The proportion of correct trials was compared to a permuted baseline of randomly drawn trials for each participant then combined over participants to create a t-statistic. D) Secondary control analysis in which we reran the classification analysis for causal choice information at the time of outcome, but only on trials where lFPC was found to correctly decode pending causal choice information. Note that this test is different from A because we allowed the classifier to create a new hyperplane separating categories for only those trials in which the lFPC decoding was “correct”. For illustration, all maps are displayed at a threshold of t(19)=2.54, p<.01 uncorrected. All effects survive small volume correction in a priori defined anatomical ROIs.

To test whether pending choice information held in lFPC was directly related to the causal choice information coded during subsequent credit assignment we used an “information connectivity” (IC) analysis, which seeks to identify how information is shared between brain regions (Coutanche & Thompson-Schill, 2013). Specifically, we tested the correlation between the fidelity of the previous choice representation when in a pending state, and the same causal choice representation during subsequent credit assignment. We began using a SVM to classify representations of the causal choice during the interim feedback period in voxels in the lFPC that were shown to code this information in our previous analysis (thresholded at *t*(19) = 2.54, p<.01).

Note that this relatively liberal threshold simply allows for the inclusion of more voxels for a statistically independent test in a left-out set of trials, thereby obviating selection bias. In a left-out set of trials, we calculated the distances between the estimated hyperplane and trial-level voxel activation patterns, and then signed these distances such that positive distances reflected “correct” classifications and negative distances reflected “incorrect” classifications. These signed distances allow us to relate both success in decoding information, as well as failures, between regions. Next, we applied the same method to quantify and sign the distances when decoding the same causal choices at the time of credit assignment – that is, when viewing the relevant outcome in the next trial. Finally, we correlated the decoding distances of causal choices in a pending state in lFPC with decoding distances of these choices during credit assignment in our lOFC and HC ROIs. This allowed us to assess whether the fidelity of pending causal choices representations in lFPC predicts the fidelity of representations during credit assignment in the lOFC and HC.

This analysis revealed strong IC between representations in lFPC at feedback on trial *t* and the representations in lOFC and HC during feedback on trial *t+1.* Specifically, we found significant correlations in decoding distance between lFPC and bilateral lOFC ([x,y,z] = [-32,24, -22], *t*(19) = 3.81, [x,y,z] = [20, 38, -14], *t*(19) = 3.87, *pTFCE* <0.05 ROI corrected]) and bilateral HC ([x,y,z] = [-28, -10, -24], *t*(19) = 3.41, [x,y,z] = [22, -10, -24], *t*(19) = 4.21, *pTFCE* <0.05 ROI corrected]), Fig. 3C). Subsequent analyses confirmed that this effect was due to these regions showing a significant increase in positive (correct) decoding in trials where pending information could be positively (correctly) decoded in lFPC, and not simply due to a reduction in incorrect information fidelity (see Fig. 3C & 3D). This finding is consistent with the coding of the causal choice during feedback in lOFC and HC being dependent on that causal choice being faithfully maintained in a pending state in the lFPC.

#### HC represents task-independent stimulus identity at feedback

Next, we tested whether the content of past choice coding at feedback includes a stimulus identity code that is reinstated during credit assignment. To test for task-independent representations of the causal stimuli, we trained a linear SVM to distinguish neural patterns evoked when participants passively viewed each shape in “template trials” (see Methods). Importantly, these were presented outside the context of the learning task and were not connected to a specific action or outcome. We then tested the classifier on neural patterns evoked at the time of feedback during the learning task. This revealed significant decoding of the causal stimulus identity at the time of feedback when averaged across direct and indirect conditions, in the left HC (Fig. 4A; [x,y,z] = [-26, -16, - 16], *t*(19) = 5.20, *pTFCE* < 0.001 ROI-corrected; right hemisphere all *pTFCE>.*1). Follow-up analyses showed a marginally significant effect in the direct transition condition alone ([x,y,z] = [-24, -16, -14], *t*(19) = 3.41, *pTFCE* = .08 ROI-corrected), and a significant effect in the indirect transition condition alone ([x,y,z] = [-28, -16, -18], *t*(19) = 3.65 *pTFCE* < 0.05). These results show that when observing an outcome, the HC reinstates task-independent representations of causal stimuli, suggesting a role for the HC in retrieving the causal stimulus identity during credit assignment.

**Figure 4.**
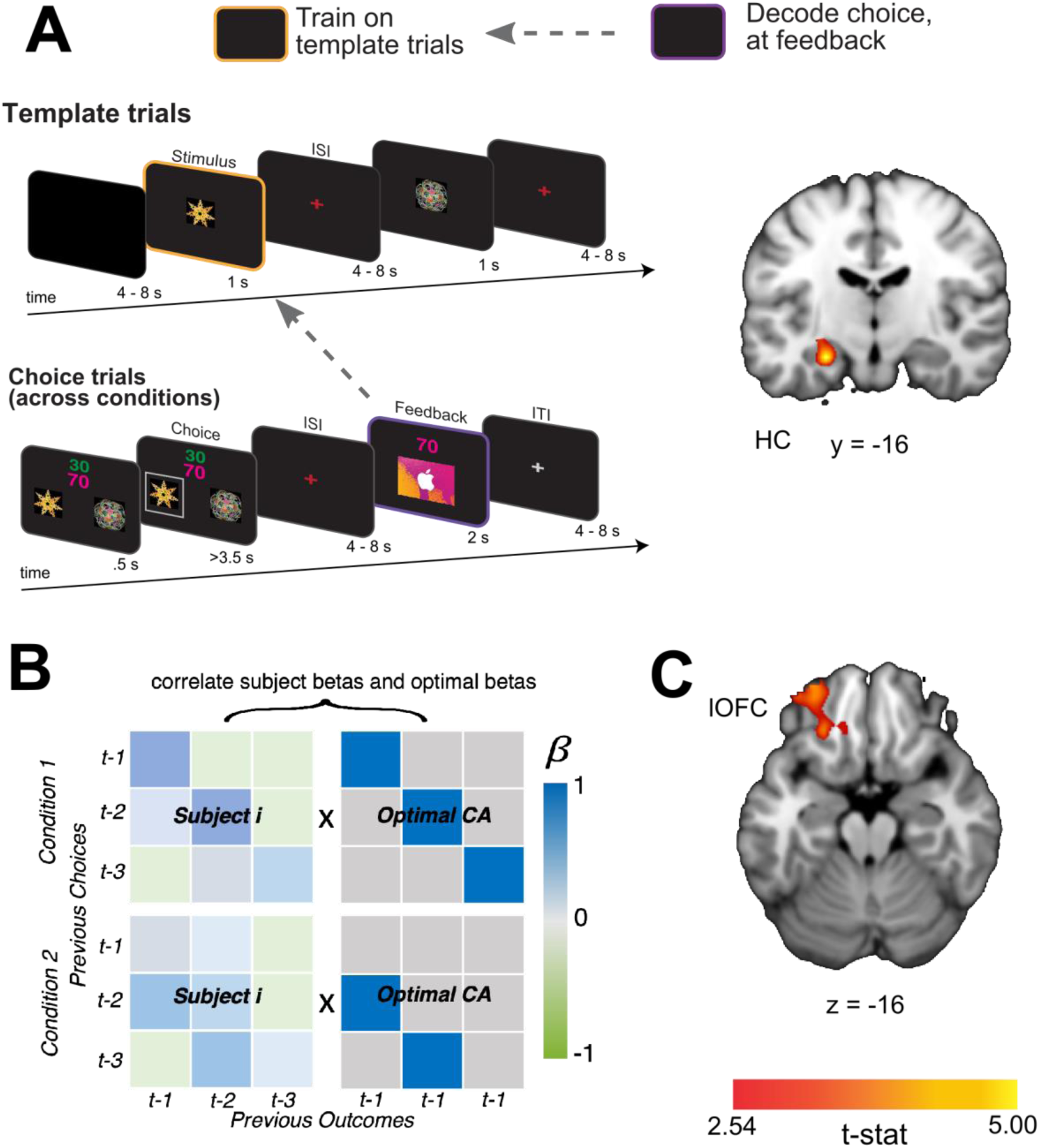
A) Schematic of the decoding procedure. In task-independent “template trials”, participants passively viewed images corresponding to the two choice stimuli and two outcome stimuli in the main task. We used these trials to train a SVM to differentiate stimuli outside the task context and then tested for representations of the causal choice stimulus at the time of feedback during the learning task. B) A coronal slice through a t-statistic map showing regions of the HC with significantly above chance decoding for the causal choice stimulus identity at the time of feedback, across conditions. In this figure, “CA” refers to “credit-assignment”. C) Analysis scheme for generating each participant’s overall credit assignment precision. β-values for each participant were taken from the behavioral model predicting current choices given all combinations of the previous three choices and outcomes (Eq.1). Each participant’s pattern of β- values (left side matrices) were correlated with a matrix representing an optimal pattern of regression betas given the task structure (right side matrices). The optimal matrix was a binary matrix with ones where credit should be assigned for a given outcomes and zeros everywhere else. D) Axial slice through a t-statistic map showing regions where decoding of the stimulus identity was significantly correlated with estimates of credit assignment precision. All maps are displayed using the same conventions as Fig. 2 and all effects survive small volume correction in a priori defined anatomical ROIs.

We reasoned further that if the HC supports credit assignment by evoking task-independent identity representations, then the extent to which this information is coded in the HC should be intimately tied to behavioral estimates of credit assignment precision. Alternatively, identity representations in the HC might support credit assignment processes in lOFC, such that the extent to which this information is represented in lOFC is predictive of precise credit assignment. To test these predictions, we estimated each participant’s overall credit assignment precision by correlating their pattern of β-values from the logistic regression models predicting choice with those of an “ideal learner” (Fig. 4B). The pattern for an ideal learner was taken to be 1 for any choice-outcome combination that reflected the true task structure, and 0 everywhere else. Higher correlations between these patterns meant that participants appropriately assigned credit to causal choices without attribution spreading to non-causal choices. We then correlated each participant’s estimated credit assignment precision with the average decoding accuracy in HC and lOFC. We found that there was a significant correlation between credit assignment precision and decoding accuracy of the causal stimulus identity reinstatement in lOFC ([x,y,z] = [-24, 34, -16], *t*(19) = 3.24, *pTFCE* <0.05 ROI-corrected), but not HC (all *pTFCE* >0.09 ROI-corrected) (Fig. 4C). These results suggest that the extent to which identity information is reinstated in lOFC is directly related to the precision with which participants link appropriate choices and outcomes together.

## Discussion

Flexible decision making in dynamic environments requires an ability to learn choice-outcome relationships across prolonged delays, which may often be punctuated by interim decisions. Understanding how the brain assigns credit for specific outcomes, and forges connections with their causal choices, is essential for models of learning and decision-making that seek to explain how organisms implement such goal-directed behaviors. The current study reveals critical roles of the lOFC and HC in such credit assignment by showing that these regions specifically represent the *causal* choice at the time the outcome is observed. Importantly, we show that when credit assignment must be delayed due to an intervening choice, representations of the causal stimulus are maintained in a “pending state” in lFPC. The fidelity of these representations determines the strength of causal choice representations in lOFC and HC when the outcome is subsequently observed. Finally, we show that the content of representations in HC includes the task-independent stimulus identities of the causal choice at the time of feedback, and the extent to which these are also represented in lOFC predicts precise credit assignment. Together, these results show that lOFC and HC adaptively use the task structure to associate identity-specific representations of causal choices to their resultant outcomes during learning and provide novel evidence for interactions between learning systems and lFPC in elaborated task structures which emulate real-world complexity.

Our finding that the lOFC instantiates a representation of the causal stimulus at the time of feedback contributes to a broader literature concerning the role of the lOFC in credit assignment. Previous research has shown that monkeys with lOFC lesions exhibit deficits in appropriately assigning credit to causal choices (Walton et al., 2010). Similarly, activity in human lOFC has been consistently associated with learning about contingencies between choices and rewards (Boorman et al., 2016; Jocham et al., 2016; Lamba et al., 2023; Noonan et al., 2017; Witkowski et al., 2022). We add to this literature by showing that the lOFC and HC contain specific multivariate patterns for inferred causal choices when an outcome is observed, suggesting that these regions are involved in updating links between choices and outcomes. Our results from the “indirect transition” condition show that these patterns are not merely representations of the most recent choice but are representations of the *causal* choice given the current task structure, and may exist alongside representations of the task structure, in the lOFC and elsewhere (Boorman et al., 2021; Park et al., 2020; Schuck et al., 2016; Seo & Lee, 2010). These findings highlight a key role for the lOFC and HC in creating links between causal states and goal-states (Boorman et al., 2021; Gardner & Schoenbaum, 2021; Howard & Kahnt, 2021; Wang & Kahnt, 2021), and suggest these regions use the specific task structure to construct causal associations between states.

While our study was designed to focus on the complexity of assigning credit in tasks with different known causal structures, another important component of real-world credit assignment is temporal ambiguity. To isolate the mechanisms which create associations between specific choices and specific outcomes, we instructed participants on the causal structure of each task, removing temporal ambiguity about the causal choice. However, our results are largely congruent with previously reported results in tasks that dissolved the typical experimental trial structure, producing temporal ambiguity, and which observed more pronounced spreading of effect, in addition to appropriate credit assignment (Jocham et al, 2016). Namely, this study found that activation in the lOFC increased only when participants received rewards contingent on a previous action, an effect that was more pronounced in subjects whose behavior reflected more accurate credit assignment. This suggests a shared lOFC mechanism for credit assignment in different types of complex environments. Whether these mechanisms extend to situations where the temporal causal structure is completely unknown remains an important question.

Importantly, we present novel evidence that representations of “pending” causal choices are stored online in the lFPC and predict the strength of causal choice representations at the time of the outcome. Our results fit precisely with theoretical proposals of lFPC functions, which propose that this region is involved in “prospective memory” and tracking alternative behaviors or task sets during ongoing behaviors which may be returned to in the future (Boorman et al., 2009; Burgess et al., 2011; Koechlin & Hyafil, 2007; Tsujimoto et al., 2011). In the “indirect transition” condition, participants needed to delay assigning credit when the first outcome was presented but return to this process when a prospective outcome was observed in the future. We show that when participants viewed outcomes for an unrelated choice, the lFPC held the content of the pending causal choice. These “pending” representations predicted the strength of subsequent causal choice representations in lOFC and HC during the next feedback period, replicating the same network we observed in the “direct transition” condition. The results extend prior work by showing that lFPC activity not only reflects statistics related to the evidence favoring pending options (Badre et al., 2009; Boorman et al., 2009, 2011; Donoso et al., 2014), but the *content* of information held in a pending state. One interpretation of these results is that the lFPC actively protects information about causal choices when potentially interfering information must be processed. Future studies will be needed to determine if the lFPC’s contributions are specific to these instances of potential interference, and whether this is a passive or active process. Nonetheless, the findings provide new evidence for the involvement of the lFPC in learning within complex task structures where the transitions between choices and outcomes are indirect - structures which abound in the real world.

Although we show evidence that lFPC is involved in maintaining specific content about causal choices during interim choices, the limited temporal resolution of fMRI makes it difficult to tell if other regions may be supporting the learning processes at timescales not detectable in the BOLD response. Thus, it is possible that the network of regions supporting credit assignment in complex tasks may be much larger. Our results provide a critical first stem in discerning the nature of interactions between cognitive subsystems that make different contributions to the learning process in these complex tasks.

A revealing aspect of our study was the inclusion of “template” trials, which allowed us to measure task-independent neural responses to the stimuli used during the learning task. By training a classifier to decode stimulus representation during passive viewing, we were able to test which regions of the brain coded the specific stimulus identity of the causal choices during credit assignment. Consistent with previous accounts of hippocampal involvement in associative learning and inference (Barron et al., 2020; Kurth-Nelson et al., 2015; Luettgau et al., 2020; Mack & Preston, 2016; Ranganath & Ritchey, 2012; Schuck & Niv, 2019; Wimmer & Shohamy, 2012), we found significant decoding of task-independent choice identities in HC across participants in both direct and indirect conditions. This suggests that the HC retrieves a representation of the stimulus identity to bind together outcomes with causal choice information at the time of credit assignment, supporting the idea that the HC is involved in linking together previous experiences of sensory information (McClelland et al., 1995). Interestingly, recent work has shown the HC neuronal ensembles code a veridical representation of stimulus identities and predicted outcomes, which are critical to inference-guided choices (Barron et al., 2020). Together, these findings imply that a state’s identity relationships constructed during credit assignment in the HC may be critical for future simulation of state-to-state transitions during outcome-guided inferences.

Interestingly, we found that the strength with which a stimulus identity can be decoded in the lOFC was correlated with behavioral measures of credit assignment, but not in HC. Recent work has shown that synchronized theta oscillations in macaques support information transfer from HC to the lOFC during value learning (Knudsen & Wallis, 2020). Disrupting these signals leads to learning deficits, suggesting that these regions work in concert to support value learning based on a relational cognitive map of the task. This synchrony between regions also finds support in human work showing strong functional connectivity and shared information between the anterior medial temporal cortex and OFC (Barnett et al., 2021; Mızrak et al., 2021; Ranganath & Ritchey, 2012). In our task, it is possible that while the HC coded task-independent identities of causal stimuli, the extent to which this information was transferred to, and represented, in the lOFC determined the efficacy of credit assignment. Future studies using methods with higher temporal resolution can elaborate on this idea by testing whether the HC and lOFC also share coherent stimulus identity information that is likewise channeled via theta phase coupling at the time of outcome, and how this information influences the credit assignment process.

In conclusion, we find that the lOFC and HC are critical to using model-based knowledge for efficiently forging links between outcomes and causal choices. Further, we show that in complex tasks where choice-outcome transitions may be interrupted, this credit assignment network relies on interactions with the lFPC, which maintains “pending” representations of causal stimuli during the interim decision. Collectively, these findings make a novel contribution to our understanding of credit assignment in the brain by illuminating the neural mechanisms which underlie linking causal choices to outcomes in complex, real-world tasks.

## Acknowledgements

Funding was provided by a Sir Henry Wellcome Postdoctoral Fellowship to EDB, a Senior Research Fellowship from the Wellcome Trust and an award from the James S. McDonnell Foundation to TEB, and a Principal Research Fellowship from the Wellcome Trust to RJD. This work was also in part supported by the Intramural Research Program at the National Institute on Drug Abuse (ZIA DA000642). The opinions expressed in this work are the authors’ own and do not reflect the view of the NIH/DHHS.

## Author Contributions

P.P.W., L.R. and E.D.B. lead the data analysis and writing of the manuscript. L.R and E.B. acquired the data. E.D.B., Z.K.-N., M.G. & T.E.B. designed and performed analyses. T.E.B. and E.D.B. conceived and designed the experiment and research question; R.J.D., T.E.B. and E.D.B. obtained funding and supervised the study.

## Data and Code Availability

Unthresholded group-level statistical maps have been deposited at NeuroVault (https://neurovault.org/collections/17702/) and are publicly available as of the date of publication. Links are listed in the key resources table. All original code has been deposited at Open Science Framework (https://osf.io/b9m6q/?view_only=eb58dd2f2076477c9bb01a8bd430b53d) and is publicly available as of the date of publication. Any additional information required to reanalyze the data reported in this paper is available from the lead contact upon request.

## Methods

### Participants

Twenty participants (11 females; 9 males; mean age = 23.5) were recruited from the general population around University College London to participate in the study. This sample size was commensurate with previous studies similar in design (Boorman et al., 2016; Howard et al., 2015; Jocham et al., 2016). Using an independent, unpublished data set, we conducted a power analysis for the desire neural effect in lOFC. We found that this number of participants had 84% power to detect this effect (Fig. S8). Participants were paid £10 and obtained a gift card of various amounts depending on their performance in the task. None of the participants reported a history of neurological or psychiatric disorder. All participants spoke fluent English and had normal or corrected-to-normal vision. The study was approved by the UCL Research Ethics Committee (Project ID Number: 3450/002), and all participants gave written informed consent.

### Task Design

#### Learning task

Participants completed a learning task in which they tracked associations between abstract shapes and specific reward identities (gift cards to two different stores), which were rated for approximately equal desirability. In each trial, participants selected one of two abstract shapes, which were randomly presented on either the left or right side of the screen. Decisions were based on two pieces of information: (1) inferred estimates of the probability that a particular shape would lead to each gift card based on the history of previous trials, and (2) the point value of each gift card on the current trial (Fig. 1A-C). Participants were informed prior to starting the task that one of the trials would be chosen at random to count “for real” at the end of the experiment. For this trial, they would receive money on the awarded gift card that was commensurate with the number of associated points (number of points divided by four). Point values for each outcome were presented as two numbers at the top of the screen, with the color of each number indicating the associated gift card identity. Their position relative to each other (top or bottom) was determined randomly on each trial.

Each shape had a specific probability of leading to each outcome and an inverse probability of leading to the other outcome. For example, shape 1 (S1) might lead to a Starbucks gift card with probability p_1_ and to an iTunes gift card with probability 1-p_1_. Shape 2 (S2) would lead to the same outcomes but with independent probabilities p_2_ and 1-p_2_, respectively. These true probabilities would drift independently over the course of the experiment, meaning that information about outcome probabilities could not be shared across shapes. On any given trial, the number of points that could be won for each gift card ranged from 20 to 100, with a minimum difference of at least 15 points. Although these magnitudes were predetermined, participants were told they were randomly generated at the beginning of each trial and that it was not useful to track them (Pearson correlation between magnitudes in trial n and n+1 was less than .2). Instead, to maximize rewards, participants had to track the probability that a shape led to each outcome and combine this with the reward magnitudes associated with each outcome on the current trial.

Each trial began with viewing the two possible choices for 0.5s, during which selection was not possible. They then had 3.5s to make their selection between the two options. The selected shape was highlighted for 0.5s, before proceeding to the interstimulus interval (ISI), which lasted for a randomly selected duration between 4s and 8s. The outcome was then presented for 2000ms before a jittered inter-trial-interval (ITI) of 4s to 8s.

Participants did not have any prior knowledge about choice-outcome associations or how quickly these associations might change, but they knew that they could change throughout the task. Therefore, participants needed to infer both the current associative contingency for each shape and when these contingencies changed from their history of choices and observed outcomes.

#### Template task

Each run of the scanning session began with a “template task”. In this task, participants passively viewed a sequence of all four stimuli (two shapes and two gift cards), individually presented in random order. To ensure that participants were paying attention during passive viewing, they were presented with 4 “catch trials” which occurred at random between images. In catch trials, all four stimuli were presented simultaneously, and participants were asked to indicate which stimulus had just been presented (Fig. S6). Participants were told they could earn an additional £10 on the selected gift card if they responded correctly. However, they would be deducted £1 for each incorrect response or for not making responses in time (max response time = 3s). Average accuracy for these catch trials was generally high (mean = .75, std=.15). Participants viewed each item for 1s followed by a 2.5s ISI.

#### Stimuli

Two visually distinct abstract shapes were used as choice objects. These shapes were randomly assigned to serve as S1 or S2 for each participant. The two gift cards were chosen to serve as reward identities during the experiment from 6 different possible gift cards (iTunes, Argos, Blackwells, Marks & Spencers, Boots, and Starbucks). Each participant rated the 6 gift cards on a scale from 0 (not preferable) to 100 (extremely preferable). The two gift cards were selected to have the minimal difference in ratings among the highest rated gift cards. This was done to prevent a strong preference for one outcome over the other. All stimuli were presented on a computer running Presentation® software (Version 18.1, www.neurobs.com).

#### Task-schedule and procedure

We generated a reward schedule that predetermined the outcome obtained for each choice on each trial, but this schedule was unknown to the participants. We optimized the schedule such that an ideal Bayesian learner (see Bayesian Computational model) would choose each shape and receive each outcome approximately an equal number of times (percent of overall trials where S1 was chosen was between 42% and 57%). This was done to reduce the potential for sampling bias in planned multivariate analyses. The schedule of outcomes for each shape was generated with independently drifting probabilities so participants could not learn anything about one shape from observing the outcome of the other shape (see Fig. S1).

Participants completed three scanning runs in one session. The first two runs began with the template task, which was followed by the learning task (37 trials of the direct transition condition, then 37 trials of the indirect transition condition). The third run consisted of only the template task. The learning task began with instructions stating, “Your latest choice”, indicating that participants were in the direct transition condition. After 37 trials, a second instruction screen showed “Your previous choice” indicating that participants were about the start indirect transition condition. Participants knew that in the indirect transition condition, the first outcome observed was not linked to any choice.

In each run, we included three “bonus trials” (two in the direct transition condition and one in the indirect transition condition), distributed throughout choice trials, which occurred between a choice and the outcome. Participants were shown the two gift cards on either side of a question mark and were given the chance to predict which outcome they would receive in the upcoming feedback period. For each correct gift card prediction, they received an additional £3 on the gift card they would receive at the end.

#### Behavioral Training

Prior to each scanning session, participants completed a shortened (76 trials) behavioral training session. In the training session, participants completed a practice version of the choice task, which had a unique reward schedule. Prior to the practice trials, participants were verbally given a “comprehension quiz” to verify they understood key elements of the task, such as the difference between choice-outcome transitions in each condition. Finally, the distribution of ISI and ITI durations for this session was constrained to 2s to 4s.

#### MRI data acquisition and preprocessing

The brain images were acquired using a 32-channel head coil from a 3 Tesla Siemens Trio scanner. We used a T2*-weighted echo-planar imaging (EPI) sequence to collect 43 2mm slices in ascending order, with 1 mm gaps. The in-plane resolution was of 3 x 3 mm, with a repetition time (TR) of 3.01s and echo-time (TE) of 70ms. We set the slice angle to a 30-degree tilt relative to the rostro-caudal axis to minimize signal loss from the lOFC (Weiskopf et al., 2006) and applied a local z-shim with a moment of -0.4 mT/m to the OFC. The first five volumes of each block were discarded to allow for T1 equilibration effects. For accurate registration of the EPI to a standard space, we acquired a T1-weighted anatomical scan with a magnetization-prepared rapid gradient echo sequence (MPRAGE) with a 1 × 1 × 1 mm resolution. Finally, to measure and correct for geometric distortions due to susceptibility-induced field inhomogeneities, a whole-brain field map with dual echo-time images (TE1 = 10 ms, TE2 = 14.76 ms, resolution 3 × 3 × 3 mm) was also acquired.

We performed slice time correction, corrected for signal bias, and realigned functional scans to the first volume in the sequence using a six-parameter rigid body transformation to correct for motion. Images were then spatially normalized by warping participant-specific images to the reference brain in the MNI (Montreal Neurological Institute) reference brain and smoothed using an 8-mm full-width at half maximum Gaussian kernel. Pre-processing was done in SPM12 (Wellcome Trust Centre for Neuroimaging, http://www.fil.ion.ucl.ac.uk/spm) using Matlab 2018a.

### Quantification and Statistical Analyses

#### Regression Analysis

To test whether participants showed a behavioral effect of learning on choice, we fit logistic regression models estimating the influence of past choice-outcome observations on choices in the current trial *t*. The regression model included the effect of the past three choices (C_t-n_) in combination with the past three observed outcomes (O_t-n_). For example, C_t-1_O_t-1_ represents the influence of the most recent choice and the most recent outcome on the current choice. The model estimates the probability of making choice C on trial *t* given all 9 combinations of previous choices and outcomes:

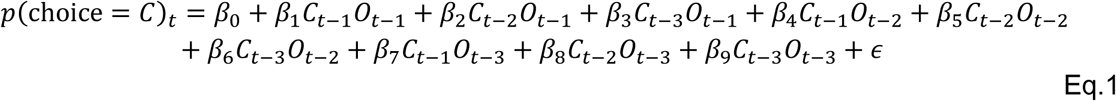

The value of C_t-n_ was taken to be 1 if they chose shape S1 on trial *t-n* and -1 if they chose S2. The value of O_t-n_ was taken to be 1 if the outcome on trial *t-n matched the currently desired outcome,* on trial *t*, and -1 if it did not. The currently desired outcome was assumed to be the outcome with the largest point value in each trial. Thus, the value of C_t-n_O_t-n_ for each trial was 1 if choice C led to the currently desired outcome *n-*trials back and -1 if it did not:

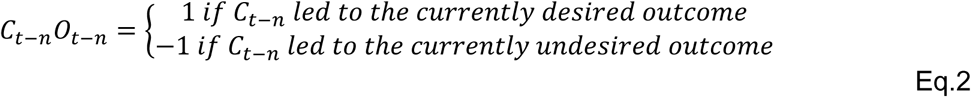

We fit separate regression models for each condition in each run for every participant. We then averaged the resulting regression coefficients (β) across runs, resulting the participant specific influence of previous decisions on the current choice.

#### Bayesian Computational model

We used a Bayesian computational model to predict choices in each trial *t* based on each participant’s previously observed shape-outcome relationships (i.e., the estimated associative probability), and reward magnitudes in the current trial. We briefly describe the model here, but a full description can be found in (Behrens et al., 2007; see also Arulampalam et al., 2002 for a related model).

Since the true probability of the associative contingencies cannot be observed, the model estimated, in a Markovian fashion, the subjective belief that making a given shape (*S*) would lead to outcome 1 (O1), and to outcome 2 (O2) with the inverse probability:

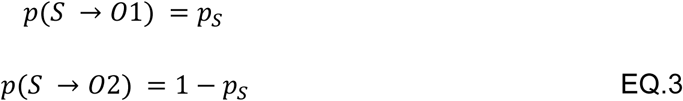

where *p_s_* denotes the associative probability of a given shape *S* leading to O1. On each trial (*t)* the model estimated the current value of *p_st_*, based on the previous observations of outcomes *y*_1:*t*_. We modeled beliefs about the likelihood of each contingency as a beta distribution over possible values of *p*_*st*_:

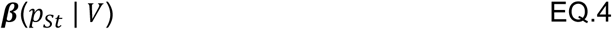

where *p*_*st*_ is the mean of the beta distribution and *V* = exp (*v*) describes the variance. A large value of *v* means that the value of *p*_*st*_ is likely to change in the next trial whereas low values of *v* mean that it is unlikely to change. Here, *v* is referred to as the “volatility” because it controls the learning rate for shape-outcome associations. The change in the estimated volatility from previous trial to the current trial is controlled by *k*. This describes the model’s belief that some level of change in the volatility is going to occur in the next trial. Because there are no constraints on values for *v*_*t*_, this distribution can be modeled as a Gaussian:

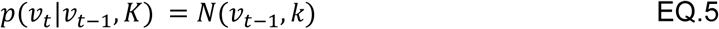

After observing each piece of evidence about the contingency between shape S and the outcome, the estimate of each parameter could then be updated following Bayes rule

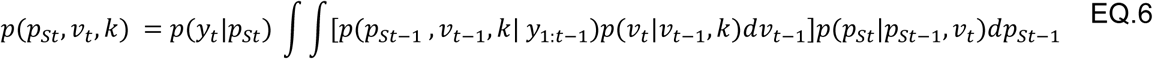

This gives us the 3-dimension joint probability of the parameters. On each trial, the learner only needs to know the estimated contingency between a shape and outcome which is performed first by marginalizing over *v* and *k*:

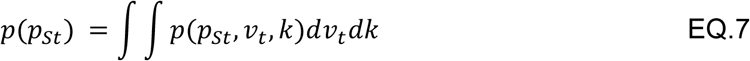

And then taking the mean of the resulting distribution.

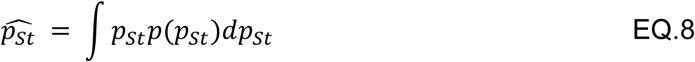

For each participant, we initialized the model with a uniform prior over the entire parameter space. All integral computations are performed using numerical grid integration. We then used the prior belief in the associative contingencies *p*^_*St*_ to compute the expected value of each shape on each trial according to the following formula:

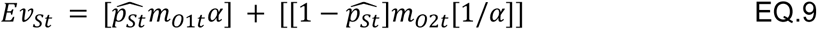

where α was a free parameter and reflected a participant’s preference for O1 over O2 (0< α <2), and *m*_o1*t*_ and *m*_o2*t*_ indicated the reward magnitudes of the outcome available in the current trial, *t*. We then measured the likelihood of each participants choice on each trial according to a SoftMax function:

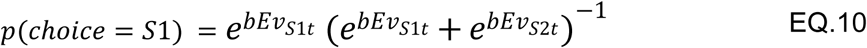

where the free parameter *b*, captured the level of sensitivity of choices to expected values (inverse temperature; 0<*b*<1). Free parameters were fitted using Markov Chain Monte Carlo (see below).

#### Value Based RL- model

This model estimated the value of each shape given the history of rewards received from choosing the shape. The value of each shape was initiated at 0, then updated using the following equation:

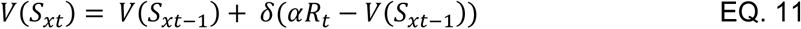

where *R*_*t*_ is the magnitude of the reward on trial *t* and α is an individual difference term estimating a participant preference for one outcome over the other (0< α <2). The learning rate (δ) was estimated for each participant to capture the magnitude of the update (0< δ <1). We entered these values into a SoftMax function to generate choice probabilities:

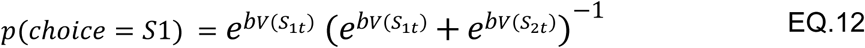

where the free parameter *b*, captured the level of sensitivity of choices to expected values (inverse temperature; 0<*b*<1). Free parameters were fitted using Markov Chain Monte Carlo (see below). Note that learning failures are not trivial to identify in our paradigm and model, because every choice is based on a participant’s preference between gift card outcomes, and the ability of the computational model to accurately estimate participants’ beliefs in the stimulus-outcome transition probabilities.

#### Parameter estimates

The Bayesian learning model has two free parameters, α and *b*. The value RL-model had an additional parameter δ. We fit these parameters independently for each participant using custom Markov Chain Monte Carlo (MCMC) code in MATLAB R2018a. Model parameters were bounded by the following: [0<α<2], [0<*b*<1], [0< δ <1] and were initialized at α=1 and *b*=.5, δ=.5. Each model was fit to maximize the likelihood of a participant’s choices given model estimates of the expected value of each choice on each trial (Eq.10; Eq.12).

#### Multivariate decoding of causal choice and pending causal choice representations

Using multivariate pattern analysis (MVPA), we aimed to identify regions of the brain that coded knowledge of causal choices during the feedback period. To test this, we estimated the BOLD activity patterns during the feedback phase for each trial using unsmoothed preprocessed images. The feedback periods were modeled as boxcars that had a constant duration lasting 2000ms from the onset of the outcome presentation in each trial. The GLM also included regressors for the decision period (modeled as boxcars with a duration equal to RT) and template presentations (modeled as boxcars with a 1000ms duration). No parametric modulators were added. Each trial was labeled according to which shape was chosen during the choice period (either S1 or S2). For our analysis of “pending” representations in the indirect transition condition, we linked these labels to the immediately following, interim feedback phase - a time when participants should be delaying credit assignment in anticipation of assigning credit in the next trial.

We used a searchlight procedure to identify regions of the brain that contained representations of the causal choice. Each searchlight consisted of a 5×5×5 voxel cube placed around a centroid voxel in the brain. Each centroid was required to have values in at least 10 of the surrounding voxels to be considered for further processing. The activity in each trial was standardized by z-scoring the β-values across voxels within each searchlight. The data were then split by blocks into training and test sets by run. We used LIBSVM (Chang & Lin, 2011) to fit linear classifiers with training data, which were subsequently used to classify data points from the test set. We iterated through this process for each of the two runs then computed the mean decoding accuracy (average proportion of correct classifications) across both classifiers. The mean decoding accuracy for each voxel was compared to a voxel-specific null distribution which was estimated by repeating this procedure while randomly assigning the labels for 100 permutations at each searchlight. The mean classification accuracy of this null distribution was subtracted off the classification accuracy of each searchlight to give us a measure of how reliably information about the causal choices could be decoded above chance. The resulting maps were then spatially smoothed using a Gaussian kernel with full width at half maximum of 8mm.

Group-level analyses were performed using a one-sample t-test on accuracy maps across participants (see *Group-level statistical inference*). We corrected for multiple comparisons over *a priori* defined ROIs in lOFC, HPC, and lFPC, and used functionally defined ROIs for lOFC in a data driven ROI analysis (see Fig. S3-5). We corrected for multiple comparisons using small volume correction TFCE. The threshold for significance remained the same in all analyses (*pTFCE* <.05).

#### Multivariate analyses of information connectivity between regions

To test whether decoding of the causal choice at feedback in the indirect transition condition depended on the strength of “pending” representations held during the interim trial, we tested whether the fidelity of representations of the pending causal choice in lFPC was associated with the fidelity of those same choices at the time of credit assignment (i.e., in the feedback phase of the next trial). We used the same decoding procedure mentioned above to classify voxel patterns at feedback in each trial, but additionally calculated the distance of each pattern from the hyperplane that divides categories. Distances were obtained using the equation specified on the LIBSVM webpage (https://www.csie.ntu.edu.tw/~cjlin/libsvm/faq.html). Patterns that are more distant from the hyperplane can be thought of as having higher fidelity, and those that are closer to the hyperplane as having less (Schuck & Niv, 2019). We then signed the distance of each point according to whether the predicted category label was correct (+ for correct, – for incorrect).

First, we calculated trial-by-trial distance from the hyperplane when causal choice information was believed to be held in a “pending” state, focusing on lFPC as our “seed-region”. For this, we calculated the average distances for voxels within the lFPC that showed significant decoding of the pending choice during the interim feedback period (*t*(19)=2.54, p<.01 uncorrected). This gave us a measure of the information about the pending item on each trial. We calculated the decoding strength of these same choices when the true outcome was shown, as a measure of the information about the causal choice during credit assignment. Here, we calculated distances for every 5×5×5 voxel cube using the same searchlight procedure we described above. Note that the decoding fidelity metric at each time point represents the decodability of *the same choice at different phases of the task.* These phases were separated by at least 10 seconds and 15 seconds on average, which can be sufficient for disentangling unique activity (Mumford et al., 2012, 2014). We then correlated the decoding distance for representations in lFPC during “pending” state and the decoding distance of those same choices at credit assignment. Thus, the correlation value between them gives us a measure of whether strong representations of pending causal choices in lFPC predict stronger representations at credit assignment.

To confirm that this correlation did not simply arise because the classifier in each region is “less wrong” when the decoder in lFPC makes correct classifications (*i.e., all classifications were wrong, but the test region was less wrong)*, we performed two control analyses. First, we calculated the frequency of correct classifications for the subset of trials in which lFPC also showed correct classifications. We then compared the frequency of correct classifications to a permuted baseline frequency by randomizing trial distances in the searchlight then recomputed the frequency of correct classifications. We subtracted the mean of the randomized baseline from the true frequency of correct classifications. This gave us a measure of decoding accuracy in each searchlight when lFPC showed correct decoding accuracy. Our second control analysis involved rerunning the classification procedure (see *Multivariate analyses of credit-assignment and pending representations),* but only for trials in which the lFPC had already shown correct decoding of the causal choice in a pending state. Again, we compared the accuracy of the classifier in each searchlight to a randomized baseline frequency by randomizing trial labels and recomputing the accuracy of the classifier. The mean of the randomized distribution was then subtracted from the classification accuracy using the true labels.

Group-level analyses were performed by Fisher-z transforming the correlation values then using a one-sample t-test on each voxel. We corrected for multiple comparisons using TFCE correction on the resulting volumes within *a priori defined* ROIs. The same thresholds were applied for group level statistical correction (*pTFCE* <.05).

#### Multivariate analyses of identity codes during credit assignment

To test whether the task-independent identity of the causal choice was reinstated during feedback, we trained a linear support vector machine (SVM) to decode representations of causal choice stimuli but trained the classifier during periods when participants passively viewed the stimuli outside of the task context (see “Template trials”). In each condition the SVM was trained on all the trials of the three template runs and tested during the feedback period of the learning task. For each participant and in each trial, we estimated the BOLD activity patterns using the same GLM as described above (see “Multivariate decoding of causal choice and pending causal choice representations”). Further, we used the same procedure in which we randomly permuted the training labels 100 times to create a null distribution of decoding accuracy. We then averaged decoding accuracy over runs and subtracted the mean of the null distribution from the true decoding accuracy of the classifier.

To test for associations between credit assignment precision and causal choice identity decoding accuracy, we first generated estimates of credit assignment precision based on each participant’s behavior during the task. For each participant we created a behavioral matrix, which included β-values from nine combinations of possible choice-outcome relationships used to assign credit when an outcome is observed (see “Regression model”). For the direct transition condition, values along the diagonal of this matrix represent appropriate credit assignment given the task structure and should have high positive values if the participant is assigning credit precisely. All other values should be near 0. A similar matrix can be generated for the indirect transition condition, but appropriate for the causal structure of this condition (see Fig. 1E). Next, we created a comparison matrix based on an idealized learner, with values of 1 in each cell that represented appropriate credit assignment for the condition, and values of 0 for non-causal relationships. We then correlated each participant specific behavioral matrix with the comparison matrix. High correlation values represent more precise credit assignment, and the average across conditions was taken to be a measure of the overall credit precision in the learning task. We then regressed each participant’s overall credit precision estimate against voxel-level decoding accuracy across participants. We corrected for multiple comparisons using TFCE correction to volumes within pre- defined ROIs. The same thresholds were applied for group-level statistical correction (*pTFCE* <.05).

#### Group-level statistical inference

Group-level testing was done using a one-sample t-test (df=19) on the cumulative functional maps generated by the first-level analysis. All first-level maps were smoothed prior to being combined and tested at the group level. To correct for multiple comparisons, we first extracted voxels from each ROI in each participant’s first-level activation map, then applied Threshold-Free Cluster Enhancement (TFCE) which uses permutation testing and accounts for both the height and extent of the cluster (Smith & Nichols, 2009). All parameters were set to default parameters (H=2, E=0.5) and used 5000 permutations for the analysis. We report effects that surpassed a *pTFCE*< .05 threshold in each ROI.

#### Region of interest selection

Regions of interest in the prefrontal cortex were generated from anatomically defined regions with unique functional connectivity fingerprints (Neubert et al., 2015). The lOFC ROIs corresponded to bilateral area BA11 (indexes 9 and 30). We included these regions because they have been previously implicated in credit assignment for causal choices, particularly in similar contingency learning tasks (Boorman et al., 2016; Jocham et al., 2016). For the lFPC, we used indexes 14 and 35. All of these ROIs were threshold at 60% inclusion criteria, although our results did not qualitatively change at different thresholds. Finally, we used *a priori* anatomically defined bilateral HC ROIs to test for effects in hippocampus (Yushkevich et al., 2015). These ROIs are illustrated in Fig. S6.

## Supplemental Figures

**Figure S1.**
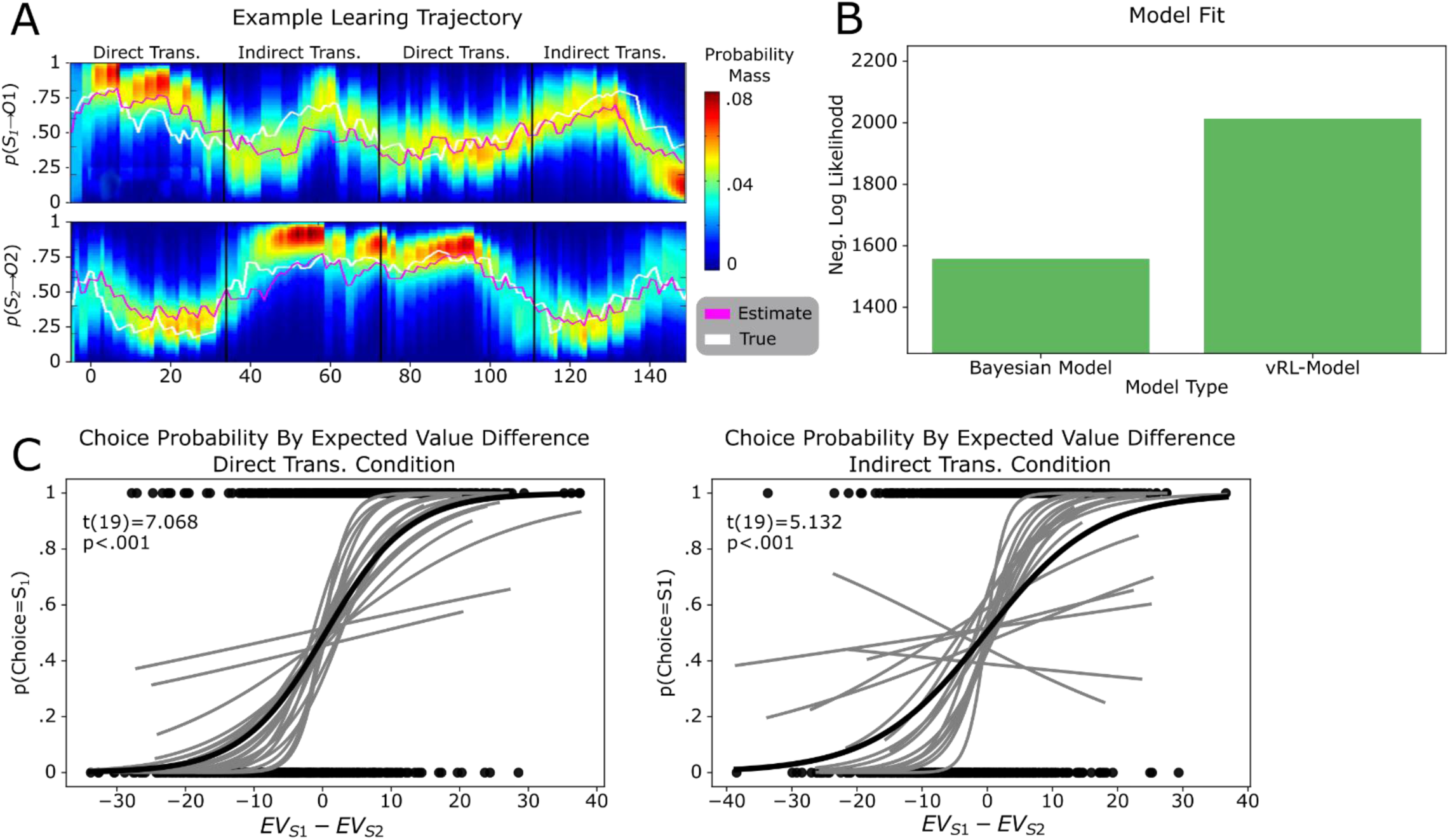
Follow up behavioral analyses. A. Example trajectory across the experiment of the belief estimates generated from the Bayesian learner. Top is the trajectory of S1, and the bottom is the trajectory of S2. While lines represent the true probability trajectory is shown in white and the estimated belief is shown in pink. Color heatmap shows the probability mass for each possible belief in S_x_ ->O1. B. Comparison of model fits between our Bayesian model and a value-based RL model (vRL) which used an interactive updating procedure to track the value of each shape based on the history of received rewards. The exceedance probability for the Bayesian model was 1, and 0 for the vRL model, suggesting that Bayesian model, which tracked transition probabilities between choices and outcomes, better fit participants actual choices compared to a value tracking model. **C**. Logistic regression curves estimating the change in choice probabilities given the expected value difference between choices. Gray line shows participant specific lines, and the black line shows the effect across groups (associated t-statistics are calculated across participants). The left side shows the effect in the direct transition condition and the right side shows the indirect transition condition.

**Figure S2.**
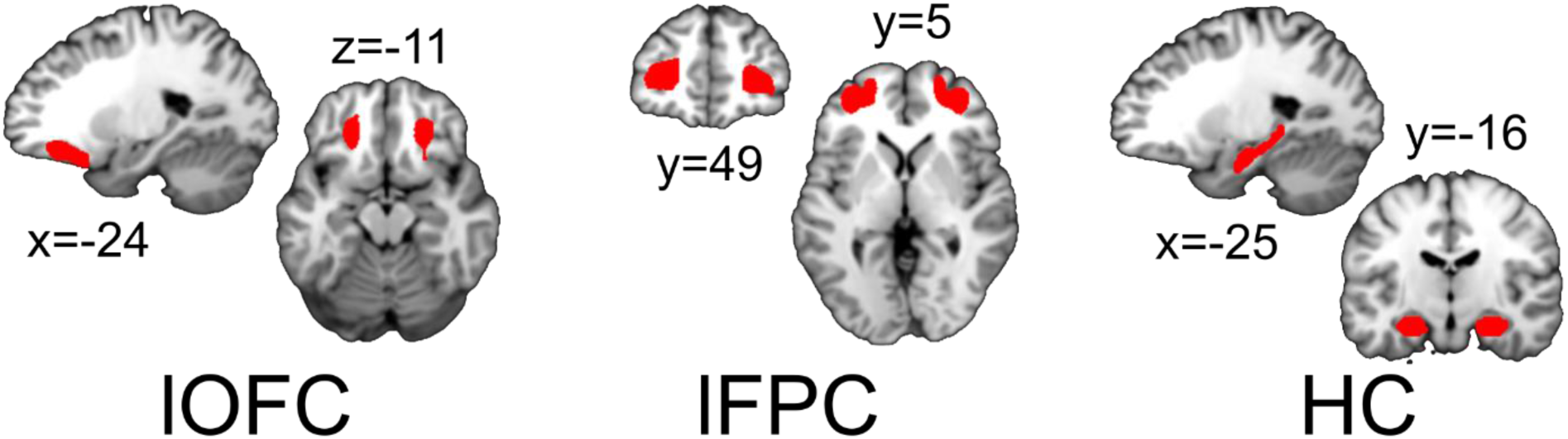
Pre-selected anatomical ROIs. Illustrations of pre-selected anatomical ROIs taken from Neubert et al, 2015. The lOFC ROI corresponds to index 9 and 30, lFPC corresponds to indexes 14 and 35. The HC ROI was defined in Yushkevich et al., 2015.

**Figure S3.**
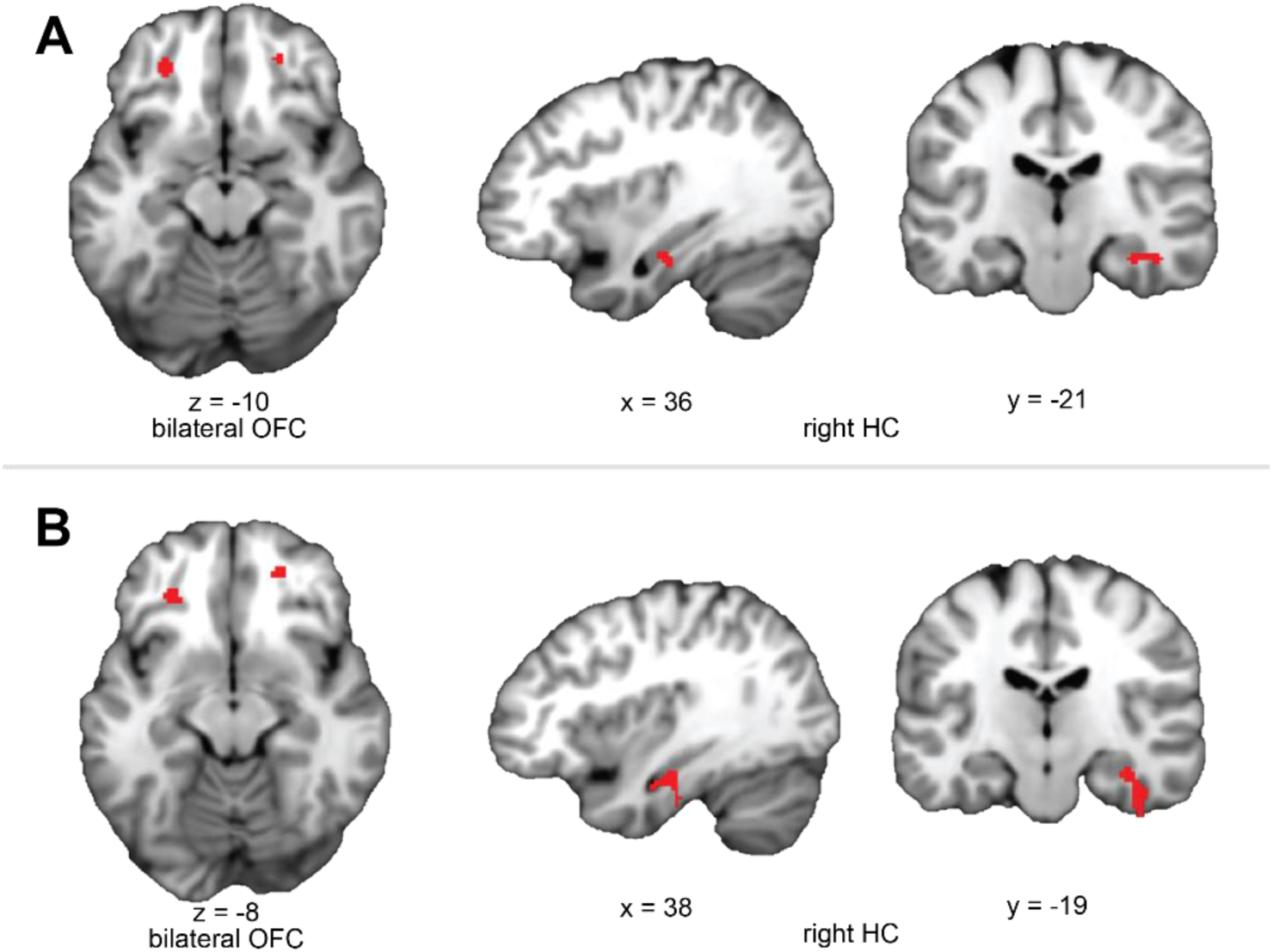
Functionally defined ROIs for in the direct transitions condition. A) Despite having *a priori* defined anatomical ROIs for our decoding analysis of the causal choice, we wanted to test whether our results depended on these ROI definitions by using a data-driven approach. Here, we trained an SVM classifier to decode representations of the causal choice in run 1 of the direct transition condition, then tested the decoder on run 2 to find regions of the orbitofrontal cortex (OFC) and hippocampus (HC) that significantly decoded causal choice representations at a significance level of *t*(19) > 2.54, p < .01, uncorrected. We then used these regions as ROIs for a separate analysis which trained the classifier in run 1 and tested the classifier in run 2. B) Shows ROIs generated from the same procedure as described in A, but the use of each run for training and testing are switched.

**Figure S4.**
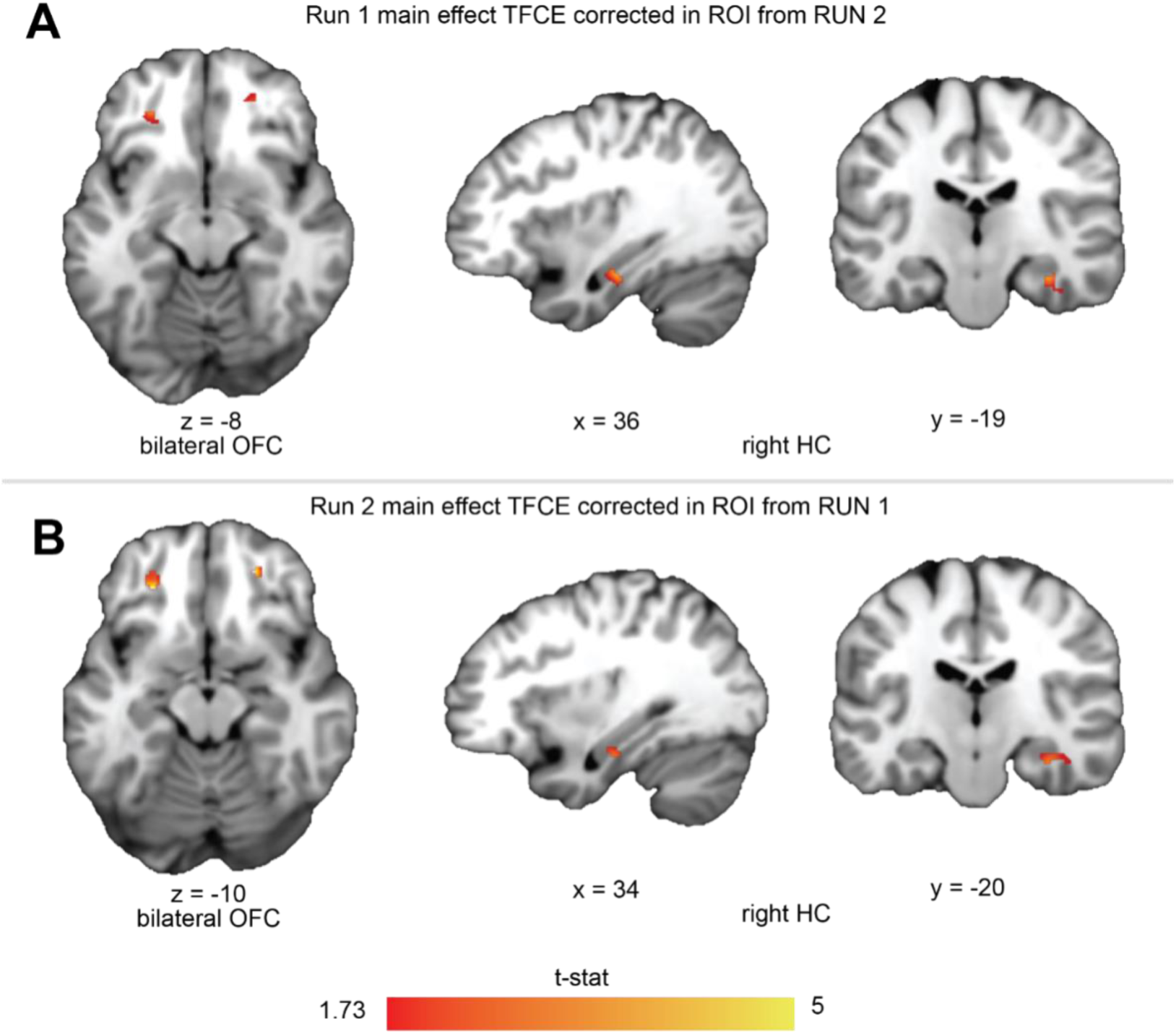
Main effect of choice decoding accuracy at the time of feedback TFCE corrected in each run of the direct transition condition. A. Regions of the OFC showing significant decoding of the causal choice in run 1 of the direct transition condition. Significance was tested using TFCE correction over voxels with the ROI generated from run 2, using the procedure described above (Fig.S1). For illustration, we show voxels that survive at threshold to t(19)=1.73, p<.05 uncorrected. B. Shows the same as A but for voxels in run 2, using the ROI generated from run 1.

**Figure S5.**
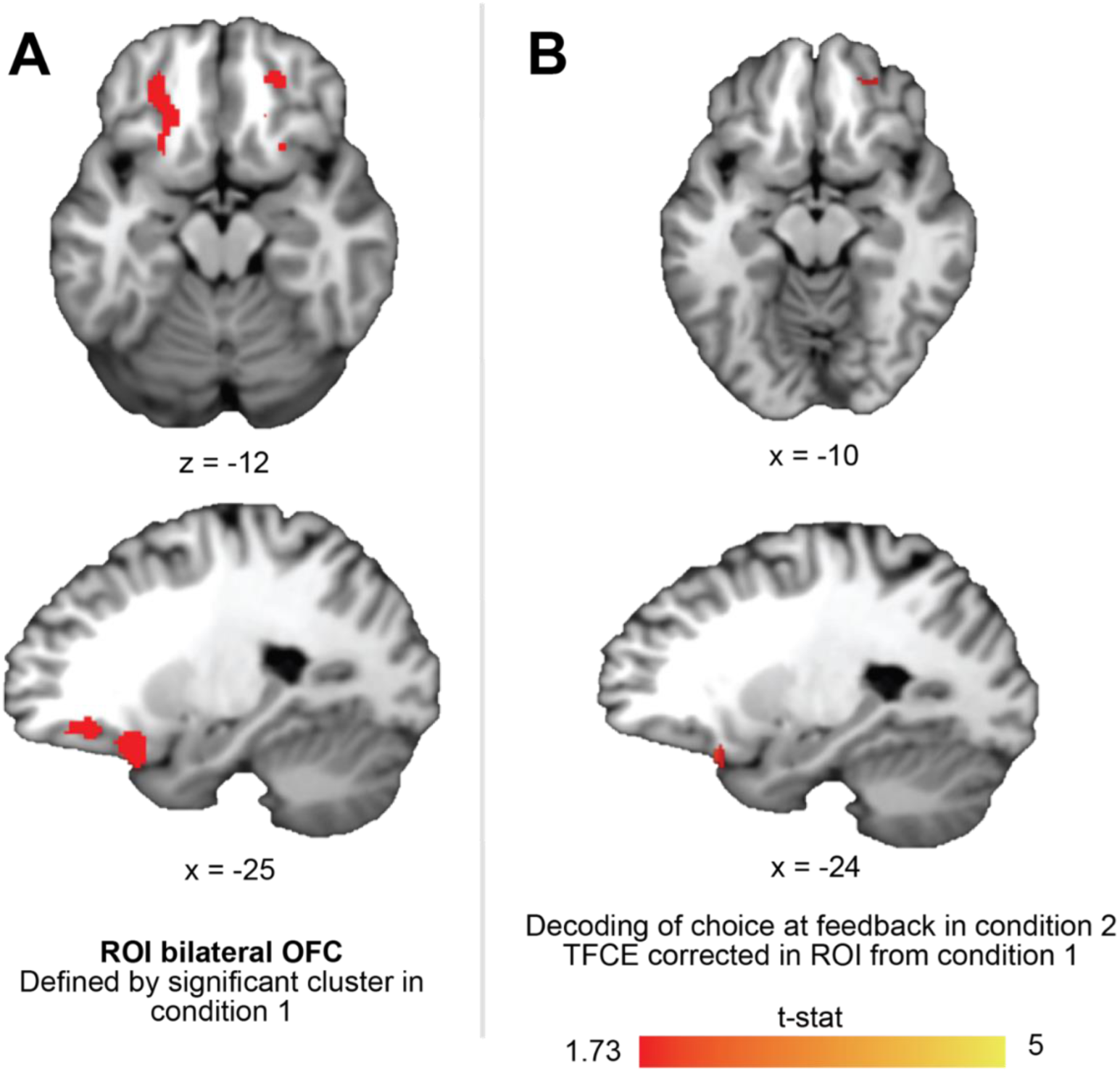
Significnant informaton connectivity between lFPC and OFC in functionally defined ROI from direct transition condition. A. We did not observe signficiant decoding of the causal choice a in bilateral OFC ROI defined by significant cluseter in in the idirected transition condition. Thus, we used the accuracy map for decoding choices at feedback during the direct transition condition (t (19) > 1.73; p < .05) in the OFC, averaged across runs. B) We then used those cluster as ROI for TFCE correction for regions of the lOFC that showed significant information connectivity with lFPC. We did this by testing for significant correlations between the trial-by-trial fidelity of pending representations in the lFPC and causal choice representation during feedback in lOFC (see Methods).

**Figure S6.**
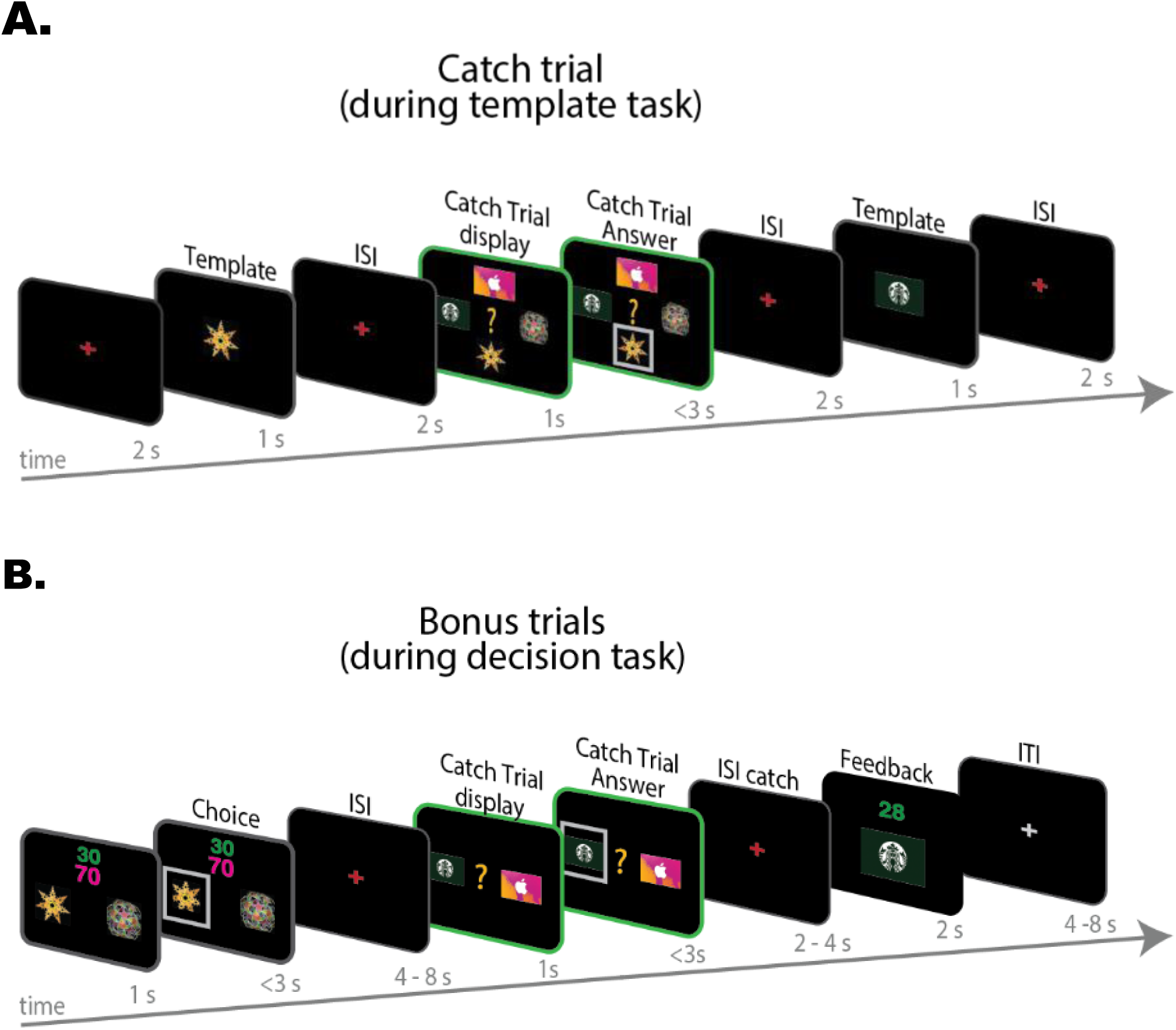
Depiction of catch trials. A. To ensure that participants where we included valuable *catch trials* in the passive observing “template task”. Participants were asked to report which image out of the four (2 gift cards and 2 stimuli) was the last one presented on the screen. They were endowed an extra £10 from which we removed £1 for every incorrect response. There were four catch trials per template run. B. The decision task included “bonus trials” in which participants could predict which gift card they expected to see on the subsequent feedback screen given their choice. They were given 3£ extra on the final gift card that was given to them for every correct answer. The first run of the direct transition condition had two catch trials; the second run had one. Both runs of the indirect transition condition had one catch trial each.

**Figure S7.**
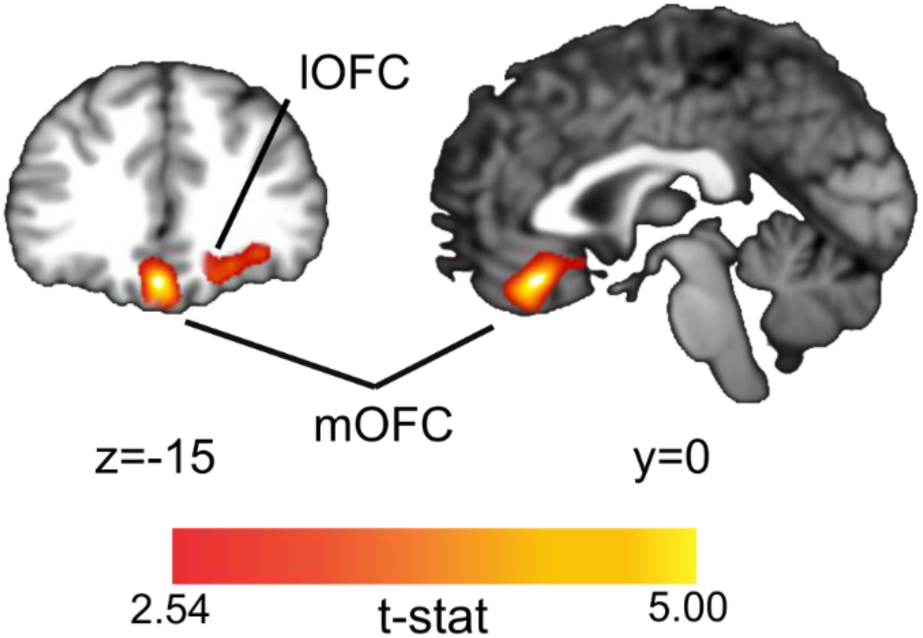
Exploratory information connectivity analysis for “Indirect transition condition”. To ascertain whether additional regions maybe involved in credit assignment beyond those that formed the focus of our study, we repeated the analysis described in Figure 2 and Figure 4 but using a whole brain search light procedure. All aspects of the analyses were the same as those previously conducted except that we corrected for multiple comparisons at the whole brain level using TFCE. For the “direct transition condition”, we found no additional regions that showed high decoding for the causal choice at the time of outcome. However, for the “indirect transition condition” we identified a region of medial OFC (mOFC) which showed information about the causal choice that was predicted by pending representation in lFPC (pTFCE<.05). The left panel shows a coronal slice through a t-statistic map, thresholded using the same conventions as Figure 4; the right panel shows a sagittal slice through the same map. These results suggest a potential role for mOFC in credit assignment uniquely during the “indirect transition condition”.

**Figure S8.**
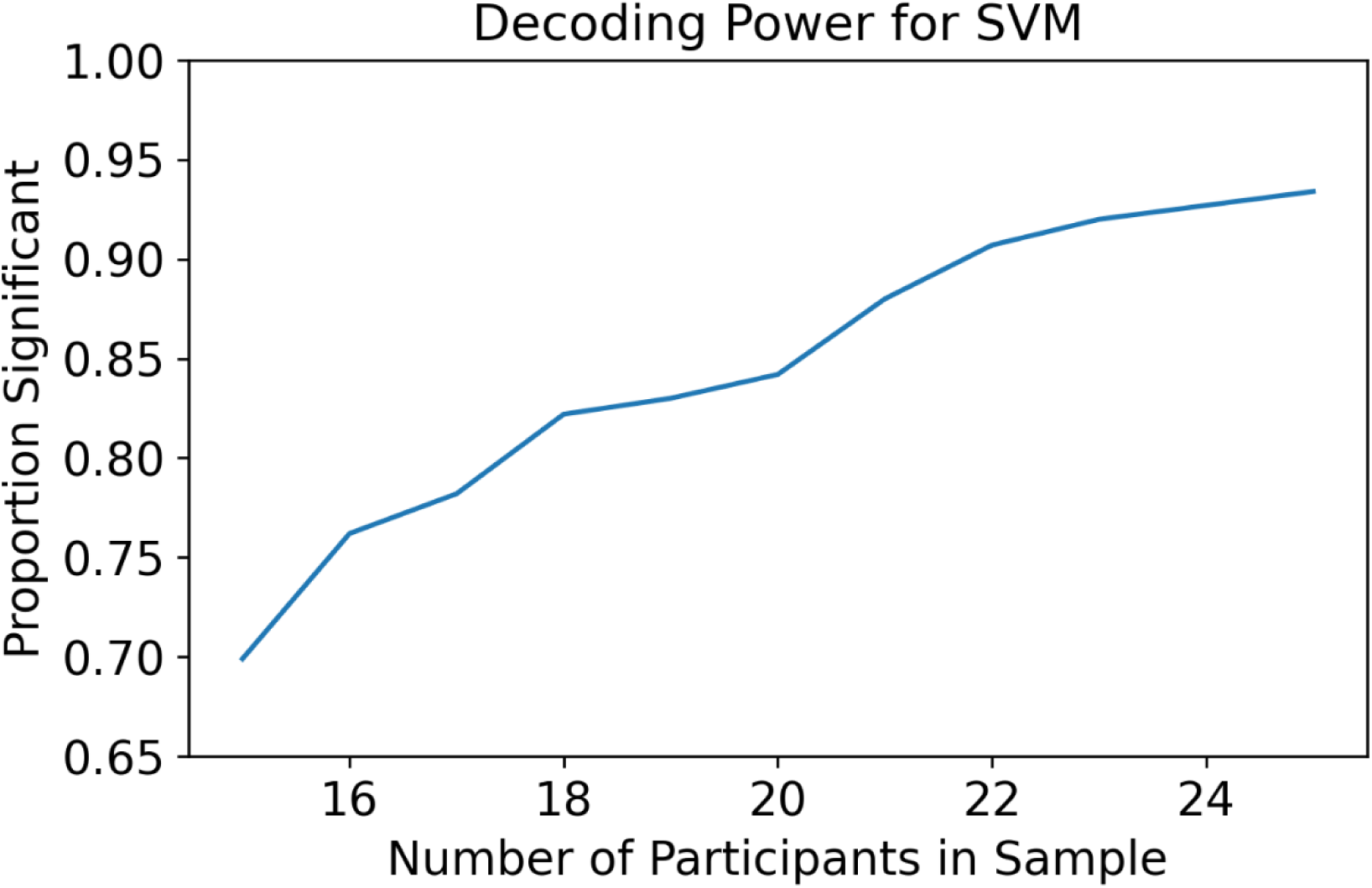
Power analysis for Reinstatement Effect in the lOFC. Power analysis using an independent data set. Twenty-eight participants competed an associative learning task, in which they learned the causal associations between four different choices, and two food rewards. We estimated voxel activity at the time of the outcome for each trial and tested for multivariate patterns of the causal choice in the lOFC, using the same procedures described in the main text (see Methods). We began by drawing 1000 samples of participants of size N, with replacement, for values of N ranging from 15 to 25. We then tested for significant decoding of the causal choice within each subset using small-volume TFCE correction. Finally, we calculated the proportion of these samples that were at or below a significance level of *pTFCE* <.05.

## References

1. Arulampalam, M. S., Maskell, S., Gordon, N., & Clapp, T. (2002). A Tutorial on Particle Filters for Online Nonlinear/Non-Gaussian Bayesian Tracking. IEEE TRANSACTIONS ON SIGNAL PROCESSING, 50(2), 723–737. 10.1109/9780470544198.ch73

2. Badre, D., Doll, B. B., Long, N. M., & Frank, M. J. (2012). Rostrolateral Prefrontal Cortex and Individual Differences in Uncertainty-Driven Exploration. Neuron, 73(3), 595–607. 10.1016/j.neuron.2011.12.025

3. Badre, D., Hoffman, J., Cooney, J. W., & D’Esposito, M. (2009). Hierarchical cognitive control deficits following damage to the human frontal lobe. Nature Neuroscience, 12(4), 515–522. 10.1038/nn.2277

4. Barbas, H., & Blatt, G. J. (1995). Topographically specific hippocampal projections target functionally distinct prefrontal areas in the rhesus monkey. Hippocampus, 5(6), 511–533. 10.1002/hipo.450050604

5. Barnett, A. J., Reilly, W., Dimsdale-Zucker, H. R., Mizrak, E., Reagh, Z., & Ranganath, C. (2021). Intrinsic connectivity reveals functionally distinct cortico-hippocampal networks in the human brain. In PLoS Biology (Vol. 19, Issue 6). 10.1371/journal.pbio.3001275

6. Barron, H. C., Reeve, H. M., Koolschijn, R. S., Perestenko, P. V., Shpektor, A., Nili, H., Rothaermel, R., Campo-Urriza, N., O’Reilly, J. X., Bannerman, D. M., Behrens, T. E. J., & Dupret, D. (2020). Neuronal Computation Underlying Inferential Reasoning in Humans and Mice. Cell, 228–243. 10.1016/j.cell.2020.08.035

7. Behrens, T. E. J., Woolrich, M. W., Walton, M. E., & Rushworth, M. F. S. (2007). Learning the value of information in an uncertain world. Nature Neuroscience, 10(9), 1214–1221. 10.1038/nn1954

8. Boorman, E. D., Behrens, T. E. J., Woolrich, M. W., & Rushworth, M. F. S. (2009). How Green Is the Grass on the Other Side? Frontopolar Cortex and the Evidence in Favor of Alternative Courses of Action. Neuron, 62(5), 733–743. 10.1016/j.neuron.2009.05.014

9. Boorman, E. D., Behrens, T. E., & Rushworth, M. F. (2011). Counterfactual Choice and Learning in a Neural Network Centered on Human Lateral Frontopolar Cortex. PLOS Biology, 9(6), e1001093. 10.1371/journal.pbio.1001093

10. Boorman, E. D., O’Doherty, J. P., Adolphs, R., & Rangel, A. (2013). The Behavioral and Neural Mechanisms Underlying the Tracking of Expertise. Neuron, 80(6), 1558–1571. 10.1016/j.neuron.2013.10.024

11. Boorman, E. D., Rajendran, V. G., O’Reilly, J. X., & Behrens, T. E. (2016). Two Anatomically and Computationally Distinct Learning Signals Predict Changes to Stimulus-Outcome Associations in Hippocampus. Neuron, 89(6), 1343–1354. 10.1016/j.neuron.2016.02.014

12. Boorman, E. D., Witkowski, P. P., Zhang, Y., & Park, S. A. (2021). The orbital frontal cortex, task structure, and inference. Behavioral Neuroscience, 135(2), 291–300. 10.1037/bne0000465

13. Burgess, P. W., Crum, J., Pinti, P., Aichelburg, C., Oliver, D., Lind, F., Power, S., Swingler, E., Hakim, U., Merla, A., Gilbert, S., Tachtsidis, I., & Hamilton, A. (2022). Prefrontal cortical activation associated with prospective memory while walking around a real- world street environment. NeuroImage, 258, 119392. 10.1016/j.neuroimage.2022.119392

14. Burgess, P. W., Dumontheil, I., & Gilbert, S. J. (2007). The gateway hypothesis of rostral prefrontal cortex (area 10) function. Trends in Cognitive Sciences, 11(7), 290–298. 10.1016/j.tics.2007.05.004

15. Burgess, P. W., Gonen-Yaacovi, G., & Volle, E. (2011). Functional neuroimaging studies of prospective memory: What have we learnt so far? Neuropsychologia, 49(8), 2246–2257. 10.1016/j.neuropsychologia.2011.02.014

16. Chang, C. C., & Lin, C. J. (2011). LIBSVM: A Library for support vector machines. ACM Transactions on Intelligent Systems and Technology, 2(3). 10.1145/1961189.1961199

17. Costa, K. M., Scholz, R., Lloyd, K., Moreno-Castilla, P., Gardner, M. P. H., Dayan, P., & Schoenbaum, G. (2023). The role of the lateral orbitofrontal cortex in creating cognitive maps. Nature Neuroscience, 26(1), 107–115. 10.1038/s41593-022-01216-0

18. Coutanche, M. N., & Thompson-Schill, S. L. (2013). Informational connectivity: Identifying synchronized discriminability of multi-voxel patterns across the brain. Frontiers in Human Neuroscience, 7. 10.3389/fnhum.2013.00015

19. Donoso, M., Collins, A. G. E., & Koechlin, E. (2014). Foundations of human reasoning in the prefrontal cortex. Science, 344(6191), 1481–1486. 10.1126/science.1252254

20. Foerde, K., & Shohamy, D. (2011). Feedback Timing Modulates Brain Systems for Learning in Humans. Journal of Neuroscience, 31(37), 13157–13167. 10.1523/JNEUROSCI.2701-11.2011

21. Gardner, M. P. H., & Schoenbaum, G. (2021). The orbitofrontal cartographer. Behavioral Neuroscience, 135(2), 267–276. 10.1037/bne0000463

22. Horner, A. J., & Burgess, N. (2013). The associative structure of memory for multi-element events. Journal of Experimental Psychology: General, 142(4), 1370–1383. 10.1037/a0033626

23. Howard, J. D., & Kahnt, T. (2021). To be specific: The role of orbitofrontal cortex in signaling reward identity. Behavioral Neuroscience, 135(2), 210–217. 10.1037/bne0000455

24. Jocham, G., Brodersen, K. H. H., Constantinescu, A. O. O., Kahn, M. C. C., Ianni, A. M., Walton, M. E. E., Rushworth, M. F. F. S., & Behrens, T. E. E. J. (2016). Reward-Guided Learning with and without Causal Attribution. Neuron, 90(1), 177–190. 10.1016/j.neuron.2016.02.018

25. Knudsen, E. B., & Wallis, J. D. (2020). Closed-Loop Theta Stimulation in the Orbitofrontal Cortex Prevents Reward-Based Learning. Neuron, 106(3), 537–547.e4. 10.1016/j.neuron.2020.02.003

26. Koechlin, E., & Hyafil, A. (2007). Anterior Prefrontal Function and the Limits of Human Decision-Making. Science, 318(5850), 594–598. 10.1126/science.1142995

27. Koechlin, E., Ody, C., & Kouneiher, F. (2003). The Architecture of Cognitive Control in the Human Prefrontal Cortex. Science, 302(5648), 1181–1185. 10.1126/science.1088545

28. Koster, R., Chadwick, M. J., Chen, Y., Berron, D., Banino, A., Düzel, E., Hassabis, D., & Kumaran, D. (2018). Big-Loop Recurrence within the Hippocampal System Supports Integration of Information across Episodes. Neuron, 99(6), 1342–1354. 10.1016/j.neuron.2018.08.009

29. Kurth-Nelson, Z., Barnes, G., Sejdinovic, D., Dolan, R., & Dayan, P. (2015). Temporal structure in associative retrieval. eLife, 4, e04919. 10.7554/eLife.04919

30. Lamba, A., Nassar, M. R., & FeldmanHall, O. (2023). Prefrontal cortex state representations shape human credit assignment. eLife, 12, e84888. 10.7554/eLife.84888

31. Luettgau, L., Tempelmann, C., Kaiser, L. F., & Jocham, G. (2020). Decisions bias future choices by modifying hippocampal associative memories. Nature Communications, 11(1), 1–14. 10.1038/s41467-020-17192-7

32. Mack, M. L., & Preston, A. R. (2016). Decisions about the past are guided by reinstatement of specific memories in the hippocampus and perirhinal cortex. NeuroImage, 127, 144–157. 10.1016/j.neuroimage.2015.12.015

33. McClelland, J. L., O’Reilly, R. C., & McNaughton, B. L. (1995). Why There Are Complementary Learning Systems in the Hippocampus and Neocortex:InsightsFrom the Successesand Failuresof Connectionist Models of Learning and Memory. Psychological Review, 102(3), 419–457.

34. Mızrak, E., Bouffard, N. R., Libby, L. A., Boorman, E. D., & Ranganath, C. (2021). The hippocampus and orbitofrontal cortex jointly represent task structure during memory- guided decision making. Cell Reports, 37(9). 10.1016/j.celrep.2021.110065

35. Mumford, J. A., Davis, T., & Poldrack, R. A. (2014). The impact of study design on pattern estimation for single-trial multivariate pattern analysis. NeuroImage. 10.1016/j.neuroimage.2014.09.026

36. Mumford, J. A., Turner, B. O., Ashby, F. G., & Poldrack, R. A. (2012). Deconvolving BOLD activation in event-related designs for multivoxel pattern classification analyses. NeuroImage. 10.1016/j.neuroimage.2011.08.076

37. Murray, E. A., & Rudebeck, P. H. (2018). Specializations for reward-guided decision-making in the primate ventral prefrontal cortex. Nature Reviews Neuroscience, 19(7), 404–417. 10.1038/s41583-018-0013-4

38. Neubert, F.-X., Mars, R. B., Sallet, J., & Rushworth, M. F. S. (2015). Connectivity reveals relationship of brain areas for reward-guided learning and decision making in human and monkey frontal cortex. Proceedings of the National Academy of Sciences, 112(20), E2695–E2704. 10.1073/pnas.1410767112

39. Noonan, M. P., Chau, B. K. H., Rushworth, M. F. S., & Fellows, L. K. (2017). Contrasting effects of medial and lateral orbitofrontal cortex lesions on credit assignment and decision-making in humans. Journal of Neuroscience, 37(29), 7023–7035. 10.1523/JNEUROSCI.0692-17.2017

40. Park, S. A., Miller, D. S., Nili, H., Ranganath, C., & Boorman, E. D. (2020). Map Making: Constructing, Combining, and Inferring on Abstract Cognitive Maps. Neuron, 107(6), 1226–1238.e8. 10.1016/j.neuron.2020.06.030

41. Ranganath, C., & Ritchey, M. (2012). Two cortical systems for memory-guided behaviour. Nature Reviews Neuroscience, 13(10), Article 10. 10.1038/nrn3338

42. Rushworth, M. F. S., Noonan, M. A. P., Boorman, E. D., Walton, M. E., & Behrens, T. E. (2011). Frontal Cortex and Reward-Guided Learning and Decision-Making. Neuron, 70(6), 1054–1069. 10.1016/j.neuron.2011.05.014

43. Schuck, N. W., Cai, M. B., Wilson, R. C., & Niv, Y. (2016). Human Orbitofrontal Cortex Represents a Cognitive Map of State Space. Neuron, 91(6), 1402–1412. 10.1016/j.neuron.2016.08.019

44. Schuck, N. W., & Niv, Y. (2019). Sequential replay of nonspatial task states in the human hippocampus. Science, 364(6447). 10.1126/science.aaw5181

45. Seo, H., & Lee, D. (2010). Orbitofrontal Cortex Assigns Credit Wisely. Neuron, 65(6), 736–738. 10.1016/j.neuron.2010.03.016

46. Shohamy, D., Myers, C. E., Hopkins, R. O., Sage, J., & Gluck, M. A. (2009). Distinct Hippocampal and Basal Ganglia Contributions to Probabilistic Learning and Reversal. Journal of Cognitive Neuroscience, 21(9), 1820–1832. 10.1162/jocn.2009.21138

47. Smith, S., & Nichols, T. E. (2009). Threshold-Free Cluster Enhancement: Addressing problems of smoothing, threshold dependence and localisation in cluster inference. Neuroimage, 44(1), 83–98.

48. Stalnaker, T. A., Cooch, N. K., & Schoenbaum, G. (2015). What the orbitofrontal cortex does not do. Nature Neuroscience, 18(5), 620–627. 10.1038/nn.3982

49. Sutton, R. S., & Barto, A. G. (2014). Reinforcement Learning: An Introduction. 10.1021/bk-1997-0674.ch022

50. Takahashi, Y. K., Roesch, M. R., Wilson, R. C., Toreson, K., O’Donnell, P., Niv, Y., & Schoenbaum, G. (2011). Expectancy-related changes in firing of dopamine neurons depend on orbitofrontal cortex. Nature Neuroscience, 14(12), 1590–1597. 10.1038/nn.2957

51. Tsujimoto, S., Genovesio, A., & Wise, S. P. (2009). Monkey orbitofrontal cortex encodes response choices near feedback time. Journal of Neuroscience, 29(8), 2569–2574. 10.1523/JNEUROSCI.5777-08.2009

52. Tsujimoto, S., Genovesio, A., & Wise, S. P. (2011). Frontal pole cortex: Encoding ends at the end of the endbrain. Trends in Cognitive Sciences, 15(4), 169–176. 10.1016/j.tics.2011.02.001

53. Walton, M. E., Behrens, T. E. J., Buckley, M. J., Rudebeck, P. H., & Rushworth, M. F. S. (2010). Separable Learning Systems in the Macaque Brain and the Role of Orbitofrontal Cortex in Contingent Learning. Neuron, 65(6), 927–939. 10.1016/j.neuron.2010.02.027

54. Wang, F., & Kahnt, T. (2021). Neural circuits for inference-based decision-making. Current Opinion in Behavioral Sciences, 41, 10–14. 10.1016/j.cobeha.2021.02.004

55. Wang, F., Schoenbaum, G., & Kahnt, T. (2020). Interactions between human orbitofrontal cortex and hippocampus support model-based inference. PLoS Biology, 18(1), 1–24. 10.1371/journal.pbio.3000578

56. Weiskopf, N., Hutton, C., Josephs, O., & Deichmann, R. (2006). Optimal EPI parameters for reduction of susceptibility-induced BOLD sensitivity losses: A whole-brain analysis at 3 T and 1.5 T. NeuroImage, 33(2), 493–504. 10.1016/j.neuroimage.2006.07.029

57. Wikenheiser, A. M., & Schoenbaum, G. (2016). Over the river, through the woods: Cognitive maps in the hippocampus and orbitofrontal cortex. Nature Reviews Neuroscience, 17(8), 513–523. 10.1038/nrn.2016.56

58. Wimmer, G. E., & Shohamy, D. (2012). Preference by Association: How Memory Mechanisms in the Hippocampus Bias Decisions. Science, 338(6104), 270–273. 10.1126/science.1223252

59. Witkowski, P. P., Park, S. A., & Boorman, E. D. (2022). Neural mechanisms of credit assignment for inferred relationships in a structured world. Neuron, 2021.12.22.473879. 10.1016/j.neuron.2022.05.021

60. Yushkevich, P. A., Amaral, R. S. C., Augustinack, J. C., Bender, A. R., Bernstein, J. D., Boccardi, M., Bocchetta, M., Burggren, A. C., Carr, V. A., Chakravarty, M. M., Chételat, G., Daugherty, A. M., Davachi, L., Ding, S.-L., Ekstrom, A., Geerlings, M. I., Hassan, A., Huang, Y., Iglesias, J. E., … Zeineh, M. M. (2015). Quantitative comparison of 21 protocols for labeling hippocampal subfields and parahippocampal subregions in in vivo MRI: Towards a harmonized segmentation protocol. NeuroImage, 111, 526–541. 10.1016/j.neuroimage.2015.01.004

61. Zajkowski, W. K., Kossut, M., & Wilson, R. C. (2017). A causal role for right frontopolar cortex in directed, but not random, exploration. eLife, 6, e27430. 10.7554/eLife.27430

62. Zeithamova, D., Dominick, A. L., & Preston, A. R. (2012). Hippocampal and Ventral Medial Prefrontal Activation during Retrieval-Mediated Learning Supports Novel Inference. Neuron, 75(1), 168–179. 10.1016/j.neuron.2012.05.010

